# On-target mutations confer resistance to WRN helicase inhibitors in Microsatellite Unstable Cancer Cells

**DOI:** 10.64898/2025.12.23.695783

**Authors:** Gabriele Picco, Yanhua Rao, Angham Al Saedi, Samantha J. Walker, Shriram Bhosle, Yang Lee, Sara F. Vieira, Maria Garcia-Casado, Gilberto Valdes Garcia, Kieron May, Francesco Sassi, Inigo Barrio Hernandez, Mamta Sharma, Cansu Dincer, Theo Bell, Anastasia Kavasakali, Ray Shenje, Cuthbert Martyr, Edward Brnardic, Nanhua Deng, Hristina Grigorova Dimitrova, Emre Karakoc, Sandeep Sundara Rajan, Nicola Chan, Emma Hitch, Katrina McCarten, Camille Fourneaux, Zoe Hewitson, Howard Lightfoot, Syd Barthorpe, James P. Phelan, Phil Landis, Brian Jones, Diana Munoz, Jay Prakash, Paul Barsanti, Josh Taygerly, Michael P. DeMartino, Emanuel Gonçalves, Andrea Bertotti, Livio Trusolino, Mike White, Geeta Sharma, Matthew A. Coelho, Jonathan Houseley, Benjamin Schwartz, Mathew J. Garnett

## Abstract

Werner helicase inhibitors (WRNi) are in clinical development for microsatellite-unstable (MSI) tumors with defective DNA mismatch repair. Here, we investigate how cancer cell evolution shapes response to WRN inhibition and informs potential resistance mechanisms. Genome-wide CRISPR screens combined with WRN knockout did not identify bypass mechanisms, underscoring WRN’s essential, non-redundant function in MSI cells. Pharmacogenomic screens identified modulators of WRNi sensitivity, including *SMARCAL1*, which links it to WRN-MSI synthetic lethality. Semi-saturation mutagenesis of WRN and prolonged drug treatment identified on-target WRN mutations driving acquired resistance to multiple WRNi in vitro and in vivo, which was mitigated by combination with standard chemotherapies. Some resistance mutations conferred broad cross-resistance, whereas others preserved sensitivity to alternative clinical-grade WRNi with distinct mechanism of action. Our findings could inform clinical trial design by suggesting the feasibility of real-time tracking of emerging resistance and enabling early therapeutic adaptations.

**Significance:** We present the first exploration of how MSI cancer cells evolve under the selective pressure of WRN helicase inhibition, providing a framework for understanding adaptive responses to this newly identified synthetic-lethal dependency. This study identifies on-target WRN mutations as key drivers of resistance in MSI cancers, supporting the use of combination strategies with other standard-of-care treatments to prevent resistance. It highlights how mutation tracking can guide therapeutic switching to clinically available WRN inhibitors with distinct mechanisms of action, thereby refining clinical development and potentially improving biomarker-informed patient outcomes.

## Introduction

WRN helicase synthetic lethality represents a novel approach for treating patients with microsatellite unstable (MSI) cancers(1–6). Recently, three WRN inhibitors have been reported, including covalent compounds (GSK_WRN3 and VVD-133214) and a non-covalent inhibitor (HRO761) (7–9), with additional molecules in development. VVD-133214 and HRO761 have entered Phase 1 clinical trials for patients with MSI cancer, either as monotherapies or in combination with chemotherapy or immunotherapy (Clinical Trial Identifiers: NCT06004245; NCT05838768). VVD-133214 was well tolerated and had anti-tumor activity in early results from these first-in-human studies(7). During revision of this manuscript, additional WRN inhibitors entered clinical evaluation, including GSK4418959(8) (NCT06710847), MOMA-341 (NCT06974110), and NDI-219216 (NCT06898450).

Despite considerable potential, as with all existing targeted therapies, understanding and addressing mechanisms of WRNi resistance will be key to maximizing clinical benefit for patients. Yet, how MSI cancer cells adapt to genetic or pharmacological WRN inactivation remains largely unexplored. The study of drug resistance can illuminate drug mechanisms of action, inform the design of second-generation inhibitors, enable rational combination therapies, and support patient stratification. Current approaches often rely on retrospective sequencing of tumor biopsies from patients who relapse on treatment. These samples are difficult to obtain and can require years to accrue in sufficient number. Moreover, interpretation of (often rare) individual variants is challenging and usually requires experimental validation to confirm causality. As a result, sequencing of tumor biopsies and ctDNA is frequently slow to yield insights and can be difficult to interpret. By prospectively modelling resistance, as done here, pre-clinical studies provide a framework to interpret complex clinical data, enabling faster and more accurate identification of mechanisms. Such approaches complement clinical analyses and accelerate the translation of findings into therapeutic strategies, avoiding delays inherent to retrospective studies and advancing patient benefit more rapidly.

The WRN dependency in MSI cancers, driven by aberrant TA dinucleotide repeat expansions that occur across the genome due to mismatch repair deficiency(3,6), represents a unique synthetic lethal mechanism in oncology. While potential escape mechanisms have been proposed (9), they remain unconfirmed given the recent development of WRN inhibitors and novel mechanisms of WRN synthetic lethality. There are at least three plausible WRNi resistance mechanisms. Little is known about the intratumor heterogeneity of TA repeat expansions associated with WRN dependency or the number of expansions or expanded sites required to confer sensitivity (3). The selection of subclones with fewer expanded TA repeats could confer resistance (9). Second, the spontaneous evolution of ‘on-target’ mutant clones that alter drug binding could confer resistance. While an engineered mutation at the binding site CYS727 has been shown to confer resistance to WRN inhibitors by preventing drug binding (10,11), no spontaneous mutations in WRN have been reported to date. In this scenario, the potential impact on the efficacy of WRN inhibitors with different mechanisms of action remains unexplored. Third, alternative or compensatory pathways, or bypass mechanisms, could overcome WRN synthetic lethality and contribute to resistance.

Early studies have provided limited insights into potential resistance mechanisms. A recent whole-genome screening study found no evidence of genes able to rescue WRN degradation (12). However, since the experiments were conducted in a single MSI cancer cell line, the findings require validation in additional models and with orthogonal systems. Inactivation of *TP53* has been reported to abrogate apoptosis induced by WRN depletion in MSI colorectal cancer cells (13). However, *TP53* did not behave as a bona fide WRN resistance gene in preclinical models where WRN was degraded or genetically inactivated (6,14) or in preclinical assessment of WRN inhibition (10,11), although another recent study suggested otherwise (15). Finally, MUS81 has been identified as an endonuclease that induces toxic chromosome breakage upon WRN loss, with its depletion significantly reducing chromosome shattering, pKAP1 signaling, and DSB formation at (TA)n repeats (3). However, its role in modulating WRN addiction or WRNi sensitivity has yet to be investigated. No additional modifiers of WRN synthetic lethality have been reported to date.

Given the recent and ongoing development of clinical WRN inhibitors, we aimed to systematically characterise how MSI cancer cells evolve under sustained genetic or pharmacological suppression of WRN, to inform the development and clinical deployment of these agents. We employed a comprehensive approach, including CRISPR-based knockout screens, semi-saturation mutagenesis, pharmacogenomic analyses, and experiments on the evolution of adaptive resistance in human MSI cancer cells. Although we identified no universal resistance mechanisms to WRN genetic or pharmacologic inactivation, we identified biologically relevant modulators of WRN inhibitor sensitivity, unveiling new players in this synthetic lethality process. Monitoring spontaneous evolution of drug resistance following pharmacological WRN inhibition revealed the emergence of on-target WRN mutations in vitro and in vivo, some of which conferred cross-resistance to multiple WRN inhibitors. These mutations could serve as biomarkers for tracking clinical resistance. Notably, resistance driven by specific variants could be overcome by clinical WRN inhibitors with distinct mechanisms of action. These findings anticipate potential clinical challenges by revealing tumor adaptation strategies, with important implications for refining clinical trial design to detect and overcome resistance, ultimately enabling the successful development of WRN-targeted medicines.

## Results

### No Evidence of Genetic Bypass of WRN Dependency in MSI Cancer Cells

Prior to the development of potent and selective WRN inhibitors, genetic approaches were used to study the effects of WRN loss in MSI cancer models (1,2). In an HCT116 CRC MSI xenograft model with doxycycline-inducible WRN knockdown (di-WRN), *WRN* inactivation caused loss of WRN expression and strong tumor suppression, as described in our published study(1). This effect was, however, transient, with regrowth after ∼25 days accompanied by WRN expression in tumors (Supplementary Figure 1A–C), indicating that tumor suppression required sustained WRN loss. Equivalent results were reported in KM12 cell xenografts expressing inducible WRN-targeting shRNAs (2). To further explore genetic escape mechanisms and corroborate these *in vivo* findings, we performed genome-wide CRISPR/Cas9 screens in inducible *WRN* knockout MSI models, expanding beyond HCT116 to include newly generated SW48 and RKO lines (Fig. 1A and Supplementary Figure 1D–F). Screen quality metrics confirmed the technical robustness of genome-wide CRISPR screens across all models (Supplementary Figure 1F–J), although the degree of selection pressure on *WRN* knockout varied (Supplementary Figure 1G and Supplementary Table 1A-C). The only hits enriched in all three models were knockouts of SAGA transcriptional coactivator complex members (*TADA1*, *SUPT3H*, *TAF6L*, and *SUPT7L*; Fig. 1B and Supplementary Figure 2A-B). However, these were not genuine bypass mechanisms, as the resistance observed in SAGA member knockouts led to inefficient WRN knockout in those specific clones (Fig. 1C and Supplementary Figure 2C-D), consistent with the SAGA complex’s crucial role in RNA polymerase II transcription (16). Reported modulators, such as *TP53* or *MUS81,* did not confer enrichment, as their targeting sgRNAs were mildly depleted (Supplementary Figure 2E). In summary, we were unable to identify individual gene knockouts capable of rescuing *WRN* gene ablation, consistent with a recent whole-genome screen (14). Although we cannot exclude the existence of alternative mechanisms, our results indicate that, *in vitro* and *in vivo,* selective pressure favours outgrowth of a subpopulation of unedited WRN-expressing cells.

**Figure 1:**
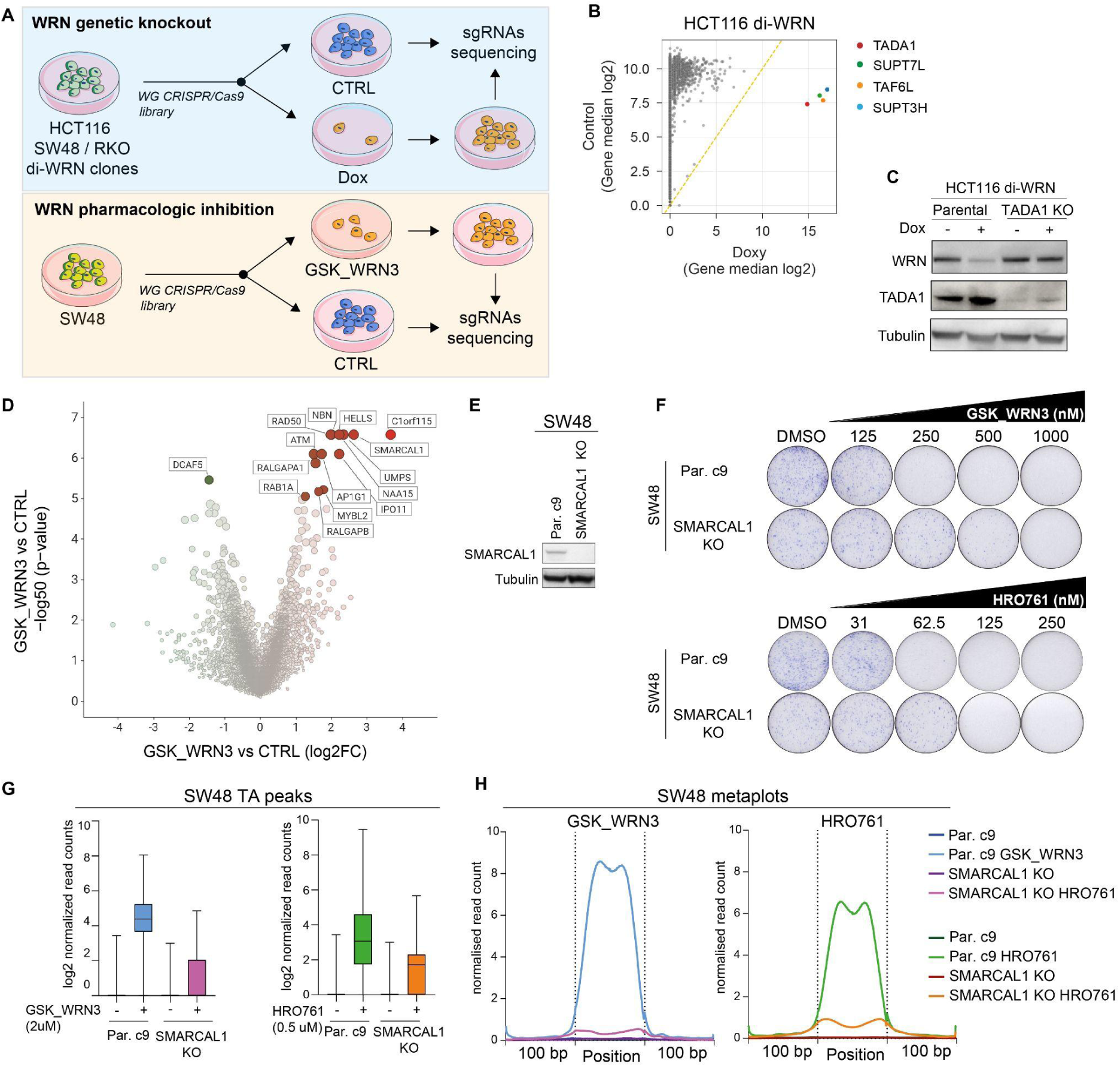
Genetic and Pharmacogenomic Screens of WRN Inhibition in MSI Cancer Cells. (A) Schematic of genetic and pharmacogenomic screening strategy. (B) Gene-level quantification of sgRNAs enriched in HCT116 WRN knockout cells (di-WRN) versus control cells. Recurrent resistance-associated genes are labelled. (C) Western blot of WRN and TADA1 in HCT116 di-WRN parental and TADA1 KO cells treated ± doxycycline. Tubulin was used as the loading control. (D) Whole-genome CRISPR/Cas9 chemogenomic dropout screen in SW48 cells treated with GSK_WRN3. The x-axis represents the log2 fold change of sgRNA representation (gene-level effect size) in WRNi-treated vs control cells, and the y-axis represents the −log10 p-value. Genes with significant negative fold changes (potential sensitisers) are highlighted in green, and those with significant positive fold changes (potential resistors) are highlighted in red. (E) Western blot for SMARCAL1 in SW48 parental cells and their corresponding SMARCAL1 knockout (KO) derivative. Tubulin is used as a loading control. (F) Clonogenic assays assessing the sensitivity of SW48 parental and SMARCAL1 KO cells to increasing concentrations of GSK_WRN3 (upper panel) and HRO761 (lower panel). Cells were treated with DMSO as a control. (G) TrAEL-seq data at TA-rich peaks in SW48 parental and SMARCAL1 KO cells treated with GSK_WRN3 (1 μM, left) or HRO761 (0.5 μM, right). (H) Metaplots of normalized TrAEL-seq read counts centred around TA dinucleotide repeats in SW48 parental and SMARCAL1 KO cells treated with GSK_WRN3 (left) or HRO761 (right).

### Pharmacogenomic Screens Identify Modulators of WRNi Sensitivity

With the advent of potent and selective WRN inhibitors, investigating modifiers of WRN pharmacological inhibition is now both possible and clinically relevant. Notably, gene disruption and pharmacological inhibition can yield distinct phenotypes, underscoring the need for direct evaluation of WRNi modifiers (15). We performed whole-genome CRISPR/Cas9-based chemogenomic screens in SW48 cells treated with a sub-lethal concentration of the covalent WRN inhibitor GSK_WRN3(10) to detect genes associated with sensitization and resistance (Fig. 1A). Performance metrics confirmed high-quality screens (Supplementary Figure 3A-D and Supplementary Figure 2).

*DCAF5* was the only significant sensitizing gene, likely due in part to the high intrinsic sensitivity of MSI cells to WRNi, reducing the ability to detect sensitizers (Fig. 1D). Neither *TP53* nor *MUS81* targeting sgRNAs were significantly modulated, consistent with genetic data from screening in di-WRN cell lines (Supplementary Table 2). Moreover, we confirmed that *MUS81* knockout did not affect sensitivity to WRN pharmacologic inhibition in isogenic lines (Supplementary Figure 3E-F). Despite there being only one gene associated with increased sensitivity to WRNi, using gene set enrichment analysis (GSEA), we found that DNA repair, G2M checkpoint, and PI3K/AKT/MTOR pathway components were significantly enriched among genes with negative fold changes (NES = −1.58, FDR q-val = 0.043; NES = −1.77, FDR q-val = 0.013; NES = −1.40, FDR q-val = 0.21, respectively) (Supplementary Figure 3G). Together, these observations suggest that these pathways may influence WRNi response in MSI cells, and their inhibition could enhance therapeutic efficacy.

Several sgRNAs were significantly enriched following WRNi treatment, indicating potential resistance genes (Fig. 1D). To aid interpretation, we grouped resistance hits by GO term enrichment, revealing associations with DNA repair, telomere maintenance, and replication stress response (Supplementary Figure 3H and Supplementary Table 2B). The top hit, *C1orf115/RDD1*, is a regulator of ABCB1 localisation and drug efflux (17), and may confer resistance by enhancing the efflux of WRN inhibitors. Several DNA double-strand break repair pathway members, including the MRN complex (*MRE11*, *RAD50*, *NBN*) and *ATM*, were significant hits, with the ATM pathway also showing enrichment by GSEA (NES = 2.25, FDR q-val = 0.001, Supplementary Table 2B). In addition, *HELLS*, a chromatin remodeler indirectly supporting double-strand break repair (18), was among the top hits. Genetic knockout of *ATM* and *NBN* in independent clones, as well as pharmacological inhibition of ATM (AZD0156), mildly reduced the sensitivity of MSI cells to WRN inhibition (Supplementary Figure 3I-L), consistent with our CRISPR screen data, although the overall effect was modest. These data suggest that ATM and NBS1 contribute to WRNi-induced cytotoxicity, likely through their roles in sensing and propagating DNA damage(19).

Among the strongest resistance genes was *SMARCAL1*, a DNA annealing helicase crucial for maintaining replication forks and responding to DNA damage (20,21). Clonogenic assays in two independent MSI cancer cell lines with *SMARCAL1* knockout confirmed a modest resistance effect, showing a slight (∼2-fold) reduction in sensitivity to both GSK_WRN3 and HRO761 (Fig. 1E-F and Supplementary Figure 3I-J). This effect was also apparent, but reduced, in shorter-term viability assays (Supplementary Figure 3M). SMARCAL1 has been recently reported to suppress DSB formation at TA-rich repeats (22,23). Thus, to gain mechanistic insights into the role of SMARCAL1 in mediating WRNi sensitivity, we compared the DSB signal at TA-repeat expansions using TrAEL-seq in *SMARCAL1* KO and parental SW48 cells after acute (24-hour) treatment with GSK_WRN3 and HRO761 (10,24). In parental lines, a strong DSB signal localised to TAs was detected after treatment with both WRN inhibitors, whereas this signal was reduced in the absence of *SMARCAL1* (Fig. 1G-H and Supplementary Figure 3N-O). This indicates a role for SMARCAL1 in mediating DSB accumulation at TA dinucleotide repeats following WRN pharmacologic inhibition and provides a rationale for the modest but reproducible resistance effect observed.

Collectively, no single gene knockout conferred robust resistance to WRN inhibition, consistent with published studies on WRN degradation(14). However, several genes did weakly modulate sensitivity to WRN inhibition, likely indirectly by influencing the DNA damage response triggered by WRNi treatment.

### Base Editing Screens Identify On-target WRN Mutations Conferring Drug Resistance

Clinical resistance to targeted therapy may arise through a selection of tumor cells with mutations in the drug target, and this can be modelled *in vitro* in cancer cells(25). Isogenic MSI cell lines engineered with alterations to the reactive cysteine (Cys727) targeted by WRN inhibitors are resistant to all WRN inhibitors to date, highlighting a possible resistance mechanism (10,11,15,26). To take an unbiased and systematic approach to identifying on-target resistance variants, we performed a base editing semi-saturation mutagenesis approach to identify residues on the WRN protein required for WRN-MSI synthetic lethal interaction (*10,25*). Clones of the colorectal KM12 and endometrial RL95-2 MSI lines, engineered to express adenine base editor (ABE) or cytidine base editor (CBE), were transduced with a library of 3,735 sgRNAs predicted to mutagenise up to ∼75% of WRN amino acids. Both lines were treated with GSK_WRN3 to assess if induced mutations modulate sensitivity to WRN inhibitors, with an additional screen in KM12 using HRO761 (Supplementary Figure 4A and Supplementary Table 3A-B). The correlation between replicate screens was high, with R values ranging from 0.79 to 0.97 across CBE and ABE screens in KM12 and RL95-2 models (Supplementary Figure 4B).

We observed minimal differential depletion of exon-targeting sgRNAs between treated and untreated conditions, indicating that no mutations strongly sensitized MSI cells to WRN inhibitors (Fig. 2A). Strikingly, sgRNAs predicted to induce mutations in Cys727 and nearby residues (723–729) were the most enriched after drug selection for both WRN inhibitors and across both models (Fig. 2A). Additionally, the WRN_GSK3 screen revealed enrichment of sgRNAs installing mutations at positions 521-522 and 852, while the HRO761 screen showed enrichment of sgRNAs targeting positions 530, 852, and 925. In a separate analysis of these data restricted to non-exonic regions, a sgRNA editing near position chr8:31,088,889 (hg38) was the sole significant resistance hit in HRO761-treated KM12 cells (Fig. 2B).

**Figure 2:**
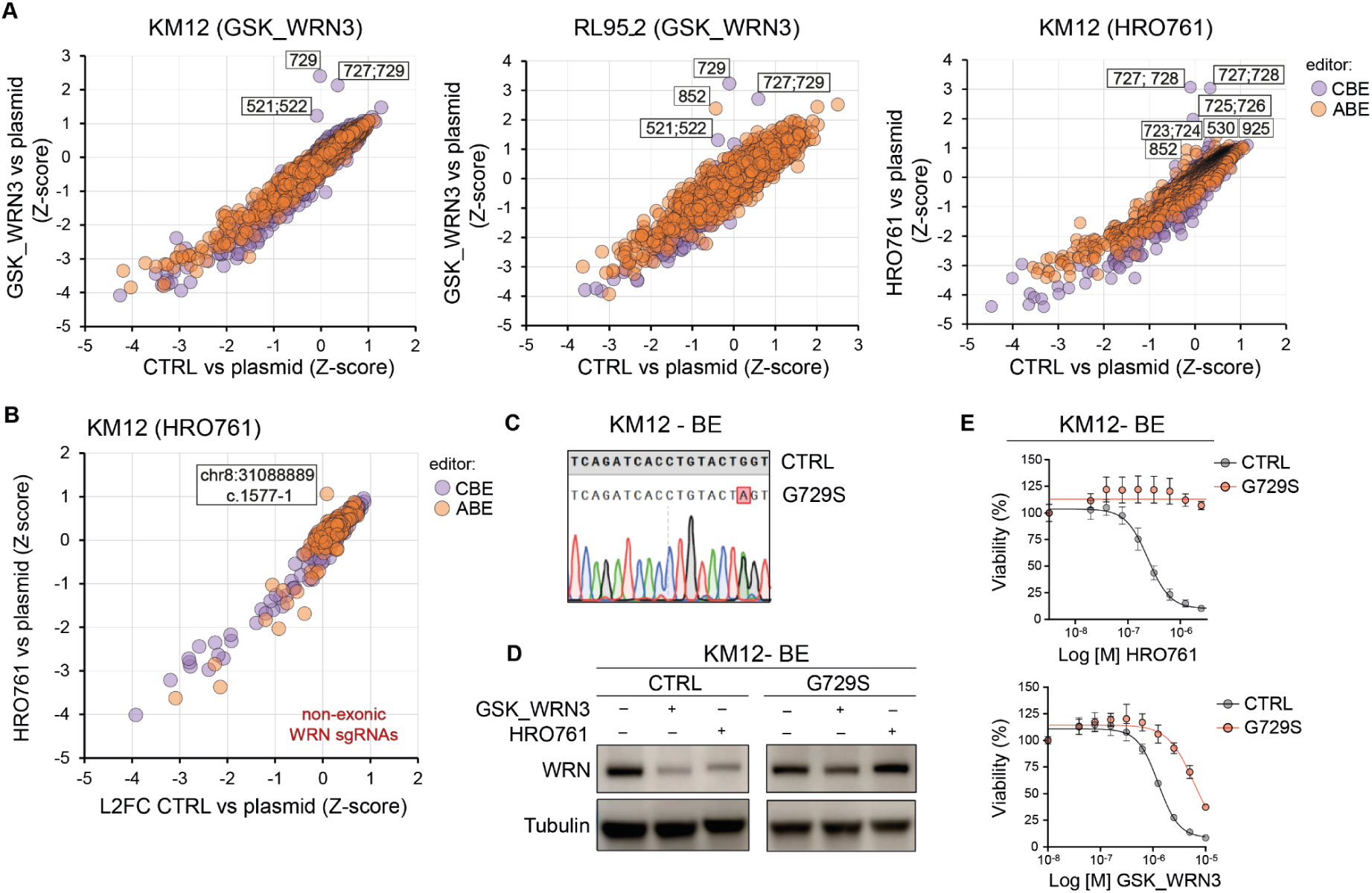
Base editing semi-saturation mutagenesis identifies on-target WRN inhibitor-resistant variants. (A) A comparison of Z-scores of exonic WRN-targeting sgRNAs between CTRL vs. plasmid and GSK_WRN3 (left and middle) or HRO761 (right) vs. plasmid in RL95-2 and KM12 cells. Points are color-coded by base editor type (CBE or ABE). (B) Scatter plot comparing Z-scores of non-exonic WRN-targeting sgRNAs between CTRL vs. plasmid and HRO761 vs. plasmid in KM12 cells. (C) Representative chromatograms of Sanger-sequenced PCR products confirming G729S mutations introduced via CRISPR-Cas9 base editing (BE) in KM12 and RL95-2 cells. (D) WRN protein levels in KM12 wild-type and G729S isogenic lines treated with either GSK_WRN3 (1 μM) or HRO761 (0.5 μM). Tubulin is used as a loading control. Representative of two independent experiments. (E) WRNi dose-response curves for KM12 wild-type and G729S isogenic lines. Data are average ± SD of three technical replicates and are representative of three independent experiments.

To validate these results, we introduced the G729S mutation via base editing (BE) in KM12 and RL95-2 BE cells (Fig. 2C and Supplementary Figure 4C). GSK_WRN3 and HRO761 failed to degrade WRN in isogenic cells carrying the G729S mutation, consistent with observed resistance (Fig. 2D-E and Supplementary Figure 4D-E).

In summary, our semi-saturation mutagenesis approach validated mutations in Cys727 and neighbouring residues that confer resistance, and identified additional critical residues, including in non-coding regions, that may be involved in resistance.

### MSI Cancer Cells Develop WRNi Resistance Through On-Target Mutations In Vitro and In Vivo

As an orthogonal method to base-editing, modeling cancer cell evolution under drug selection pressure is a powerful approach to studying mechanisms of secondary resistance, enabling identification of both on- and off-target resistance mechanisms (27). We performed an *in vitro* time-to-progression (TTP) assay, selecting SW48 colorectal cancer cell lines following continuous exposure to GSK_WRN3 and HRO761 to model the evolution of resistance to WRN inhibition in MSI cancer cells. SW48 acquired resistance to both WRN inhibitors, although with different kinetics (Fig. 3A-C and Supplementary Figure 5A). Similar patterns were observed in additional cancer cell lines from colorectal (KM12), endometrial (RL95-2, Ishikawa(Heraklio)02ER), stomach (IM-95) and ovarian (IGROV-1) cancers (Supplementary Figure 5B), indicating that MSI cancer cell lines, regardless of their tissue of origin, possess an intrinsic ability to evolve resistance to WRN pharmacologic inhibition. Building on previous evidence that irinotecan synergizes with WRN inhibitors to suppress MSI cancer growth (11,15), we performed TTP assays combining HRO761 with SN38, irinotecan’s active metabolite, at concentrations within the range of clinically achievable free drug exposure, in two MSI cell lines. Sub-lethal concentrations of SN38 synergized with HRO761 to delay growth and the onset of acquired resistance (Supplementary Figure 5C-D). Notably, in both models, combination treatment ultimately prevented the emergence of resistance for up to 60 days, outperforming either monotherapy.

**Figure 3:**
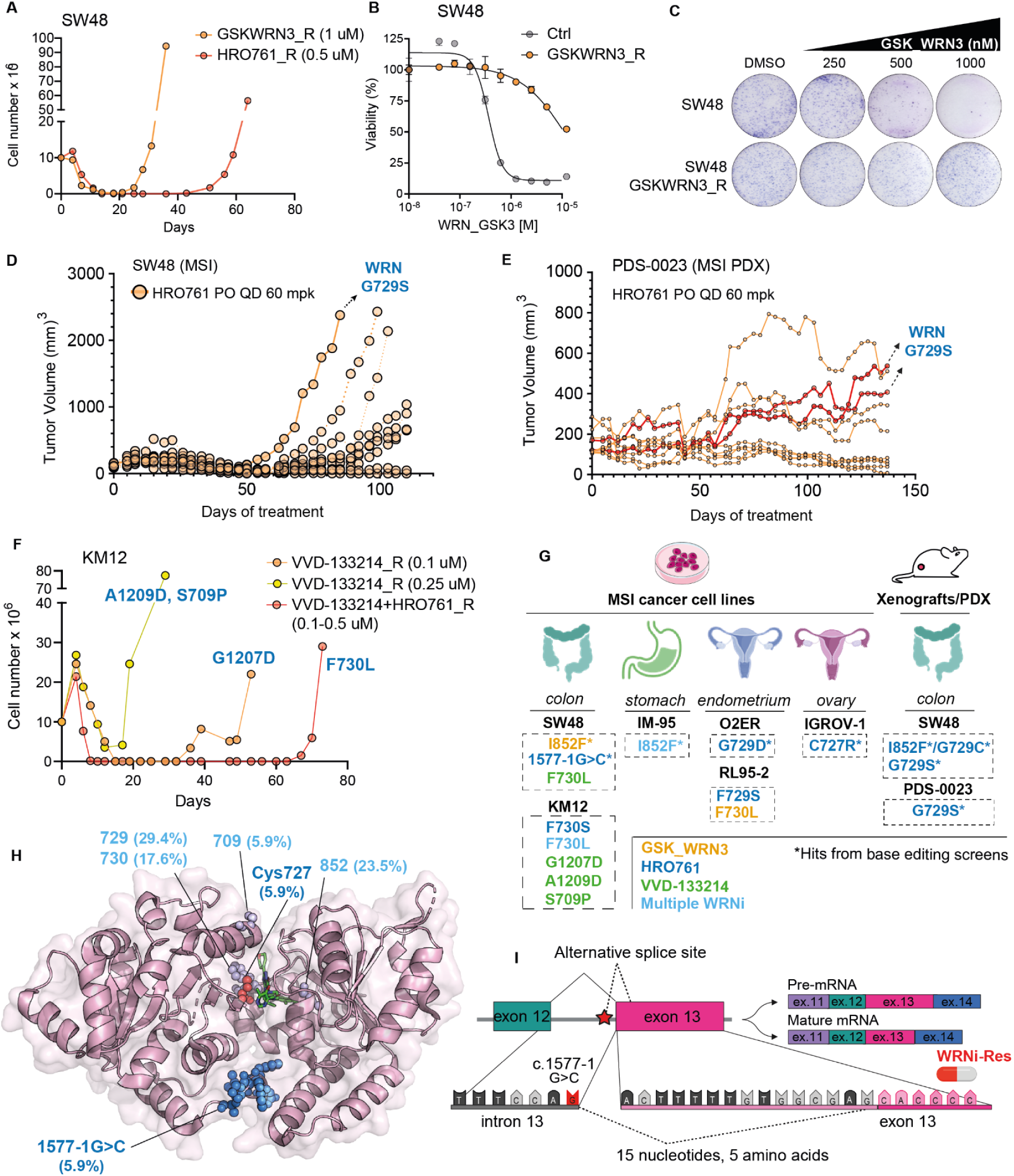
Spontaneous Emergence of WRN Mutations Drives WRNi Resistance in MSI Preclinical Models In Vitro and In Vivo. (A) TTP assays in SW48 MSI cancer cells exposed to GSK_WRN3 or HRO761 at the indicated concentrations (in parentheses). (B) Sensitivity of GSK_WRN3-resistant SW48 cells versus control cells. (C) Colony formation assay comparing differential GSK_WRN3 sensitivity of parental versus GSK_WRN3-resistant SW48 cells. (D) Individual tumor volume traces from SW48 xenografts treated with HRO761 (60 mg/kg). A subset of tumors acquired resistance, resulting in subsequent regrowth. Post-explant deep-targeted sequencing of one tumor identified WRN mutations, consistent with a role in resistance. (E) MSI PDX (PDS-0023) tumor volume measurements from mice treated with HRO761 (60 mg/kg). Two tumors with an attenuated response and early signs of resistance (highlighted in red) were selected for targeted sequencing, which revealed the G729S mutation. (F) TTP assay in KM12 colorectal cancer cells treated with indicated drug(s) and concentrations. WRN mutations identified by WGS are indicated for each model. (G) Summary of WRN inhibitor resistance mutations identified across MSI cancer cell lines, as well as cell- and patient-derived xenografts. The Ishikawa(Heraklio)02ER cell line is abbreviated as 02ER. Mutations identified in base editing screens are marked with an asterisk. The I852F and G729C mutations were detected in the same xenograft and are therefore shown linked as ‘I852F/G729C. (H) Mapping of WRN resistance mutations onto the 3D structure of the WRN ATPase domain bound to HRO761 (pdb 8PFO). The percentage of each variant across the 17 resistant models is indicated in parentheses. For the c.1577-1G>C splice site mutation, missing exon 13 residues are in blue. (I) WRN gene schematic showing the exon-truncating splice site mutation (1577-1G>C) identified in HRO761-resistant SW48 cells.

WRNi-resistant cells were then analyzed using whole-genome sequencing (WGS) to interrogate underlying mechanisms. We found that WRNi-resistant cells maintained a similar mutational load without significant changes in global copy number (Jaccard Index range: 0.77 to 0.93 (Supplementary Table 4). However, several hundred to thousands of novel mutations were detected in resistant cells, likely reflecting the expansion of subclonal populations and the high mutational load characteristic of MSI cells. We inferred the length of (TA)n repeats expansions in WRNi-resistant cells and found no difference compared to the parental unselected counterparts (Supplementary Figure 5E and Supplementary Table 4), suggesting that heterogeneity in (TA)n repeats was not a driver of acquired resistance. Moreover, no MMR gene mutations showed a substantial decrease in VAF (ΔVAF < –0.2) in resistant models compared to the parental, ruling out a reversion event leading to MMR restoration as the primary resistance mechanism (9).

Notably, sensitivity to WRN genetic inactivation by RNAi persisted in TTP WRNi-resistant SW48 cells, indicating a continued reliance on WRN protein (Supplementary Figure 5F). This suggests that WRNi has lost its ability to inhibit WRN function and suppress the growth of cancer cells. Indeed, WRNi-mediated downregulation of WRN protein levels was attenuated, and there was an abrogation of DNA damage induction, consistent with reduced inhibitor binding leading to decreased cytotoxicity (Supplementary Figure 5G). Mining WGS data, we identified a missense mutation (I852F) in the WRN gene in GSK_WRN3-resistant SW48 cells and an exon-truncating splice site mutation (1577-1G>C) in HRO761-resistant SW48 cells, both of which were absent in the parental line. Additionally, recurrent somatic WRN mutations, including I852F, C727R, G729D, F730L, F730S, and G729S, were identified in all other resistant cell models, which spontaneously emerged under the selective pressure of GSK_WRN3 and HRO761. Many of the mutations are located at or adjacent to the reactive cysteine at position 727, which is involved in drug binding, and were orthogonally validated by base editing (Fig. 2). Despite thousands of sample-specific mutations detected in resistant cells, WRN was the only gene that was recurrently mutated across all cell lines. These results provide a framework for interpreting emerging clinical data on WRN inhibitor resistance.

We additionally assessed resistance mechanisms to prolonged drug treatment *in vivo*. SW48 MSI tumor xenografts were treated long-term with two concentrations of HRO761 (60 mg/kg and 120 mg/kg), with 10 mice per group (Supplementary Figure 5H). After initial robust regression, over a longer duration (>50 days) tumor regrowth occurred in 7/10 mice at the lower dose and 1/10 at the higher dose (Fig. 3D and Supplementary Figure 5I). Based on available material, we were able to sequence one relapsed tumor per group. At 120 mg/kg, WRN mutations I852F and G729C were identified; at 60 mg/kg, I852F and G729S were detected—matching mutations found *in vitro* (Fig. 3D and Supplementary Figure 5I). In one resistant xenograft (Supplementary Figure 5I), two distinct WRN mutations (G729C and I852F) emerged in parallel in different subclones, underscoring the genetic heterogeneity and adaptability of MSI tumors under WRNi pressure. We also examined the emergence of WRN mutations in a patient-derived xenograft (PDX) model treated with HRO761. The original tumor harbored a *BRAF* V600E mutation, *MSH3* K383Rfs*32, and showed loss of *MLH1* and *PMS2*. Despite a marked response to treatment (Supplementary Figure 5J), consistent with previous findings (10,11,15), a subset of mice had attenuated responses, with tumor size stabilizing at higher volumes (Fig. 3E). Sequencing of two tumors from HRO761–treated mice revealed G729S WRN mutations, which were absent in the tumor control arm, consistent with resistance-associated clonal emergence.

To assess whether resistance consistently emerges across WRN inhibitors, we performed additional *in vitro* TTP assays with VVD-133214, administered alone or in combination with HRO761 in KM12 MSI colorectal cancer cells, and as monotherapy in SW48 cells (Fig. 3F and Supplementary Figure 5K). VVD-133214 monotherapy led to resistance in both lines, and this resistance also emerged with the combination, albeit modestly delayed compared to monotherapy. Tumor DNA sequencing identified on-target WRN mutations in both settings. G1207D and A1209D emerged with VVD-133214 monotherapy, whereas F730L also appeared under combination treatment, suggesting it may confer cross-resistance to both agents (Fig. 3F).

Overall, on-target mutations in resistant models converged on recurrent residues within the WRN helicase domain, many of which were also identified in orthogonal base editing screens (Fig. 3G and Supplementary Table 5). Mapping resistance variants onto the 3D structure of the WRN helicase domain revealed several clusters at or near cysteine 727, critical for drug binding, consistent with a mechanism of resistance through impaired inhibitor engagement (Fig. 3H). However, non-canonical mutations were also identified. The exon-truncating splice site mutation (1577-1G>C; chr8:31088889) generates a cryptic splice site, causing partial truncation of exon 13 up to position 31088904, producing an aberrant WRN transcript, which is predicted to affect the encoded protein (Fig. 3I). This finding confirms our base editing results, which identified an sgRNA targeting this non-exonic region as a resistance hit, and shows that engineering the splicing variant is sufficient to confer resistance to HRO761, consistent with its spontaneous emergence in resistant cells.

These results indicate that MSI cells from different lineages follow a consistent evolutionary trajectory both in vitro and in vivo, developing resistance to multiple clinical-grade WRN inhibitors through recurrent, on-target mutations in the WRN gene.

### MSI Cells with WRN Resistance Mutations Remain Sensitive to Clinical-Grade WRN Inhibitors with Alternative Mechanisms of Action

HRO761 and VVD-133214 are among the first WRN inhibitors to enter clinical development and inhibit WRN’s helicase enzymatic activity via distinct mechanisms: HRO761 stabilizes WRN in an extended conformation, whereas VVD-133214 irreversibly locks WRN in a compact, inactive state, impairing ATPase function (15) (Supplementary Figure 6A). To interpret how resistance variants affect inhibitor binding, we performed structural modeling. The WRN G729D mutation, located in the hinge region (residues 728–732), substitutes a flexible glycine with a bulkier, negatively charged aspartate, disrupting hinge dynamics. Energy minimization revealed local residue rearrangements that likely alter binding pocket architecture, predicted to impair the binding of both WRN inhibitors (Supplementary Figure 6B). In contrast, the recurrent I852F mutation introduces a phenylalanine side chain that protrudes into the HRO761 binding pocket, narrowing the hydrophobic cage and likely reducing drug binding, while having minimal impact on VVD-133214 (Fig. 4A). These models suggest that G729D and I852F confer distinct patterns of cross-resistance.

**Figure 4:**
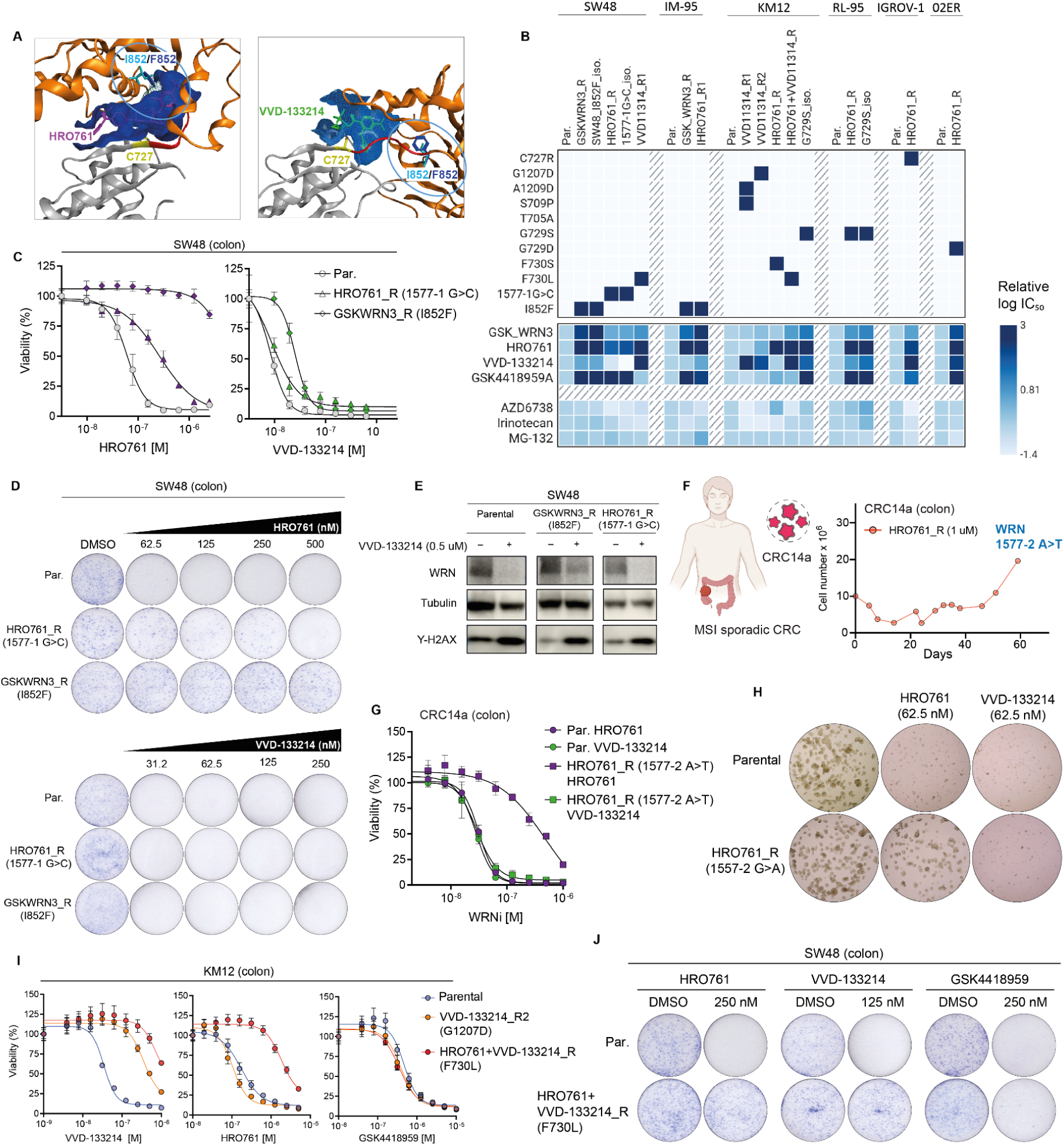
MSI Cells with WRN Resistance Mutations Retain Sensitivity to Alternative WRN Inhibitors. (A) Structural representations of HRO761 and VVD-133214 bound to WRN helicase domain. Pocket surfaces are shown for wild-type WRN (cyan) and the I852F mutant (blue) in complex with each inhibitor. (B) Heatmap of cell viability from drug screening in WRNi-resistant MSI (generated in Time-to-Progression assays) and SW48 isogenic cell lines carrying WRN mutations in heterozygosity, treated with the indicated drugs. For each cell line, log-transformed IC₅₀ values were normalized to the parental line and capped at a maximum of 3 for visualization. Cell line 02ER is an abbreviation of Ishikawa (Heraklio) 02ER. (C and D) Sensitivity of SW48 parental and WRNi-resistant cell lines (I852F, 1577-1G>C) treated with HRO761 or VVD-133214 in (C) viability or (D) clonogenic assays. (E) Immunoblot of WRN, γ-H2AX, and tubulin in SW48 parental and WRNi-resistant cells following treatment with VVD-133214. (F) Time course of CRC14a patient-derived organoid (PDO) growth following HRO761 exposure. The 1577-2 A>T variant was identified in resistant cells at day 60. (G) Sensitivity of parental and WRNi-resistant (1577-2 A>T) CRC14a PDO cells treated with HRO761 or VVD-133214. (H) Brightfield images of CRC14a PDOs after 7-day treatment with HRO761 or VVD-133214 (62.5 nM). (I) Sensitivity of KM12 parental and WRNi-resistant cells (G1207D, F730L) treated with HRO761, VVD-133214, or GSK4418959. (J) Clonogenic assays of SW48 parental and WRNi-resistant cells (F730L) treated with HRO761, VVD-133214, or GSK4418959.

To directly test this hypothesis, and more broadly compare drug cross-resistance, we profiled parental MSI cells, WRNi-resistant derivatives, and isogenic lines carrying resistance-associated WRN mutations for their sensitivity to GSK_WRN3, HRO761, and VVD-133214 (Fig. 4B and Supplementary Figure 6C). In addition, we tested the ATR inhibitor AZD6738, a recently proposed MSI-selective targeted therapy (28), and the chemotherapeutic irinotecan, which is synergistic with WRNi in MSI cancer preclinical models (11,15). In total, we evaluated over 150 different experimental conditions.

Consistent with our structure-based predictions, HRO761-resistant cells harboring a G729D mutation were cross-resistant to all three WRN inhibitors, including VVD-133214. The C727R mutation also conferred resistance to all compounds (Fig. 4B). Notably, HRO761- and GSK_WRN3-resistant cells with an I852F mutation retained sensitivity to VVD-133214, consistent with our structural predictions, and a finding confirmed in a heterozygous SW48 I852F isogenic cell line (Fig. 4B–D and Supplementary Figure 6D-E). Similarly, HRO761-resistant cell lines with splice-site mutation 1577-1G>C, as well as a heterozygous SW48 isogenic line carrying the same alteration, remained exquisitely sensitive to VVD-133214, suggesting that this alteration may not impair VVD-133214 binding or activity. Consistent with our viability data, VVD-133214 led to WRN degradation and induction of DNA damage in resistance models with an I852F or 1577-1G>C mutation, but not in a G729D mutation (Fig. 4E and Supplementary Figure 6F). The sensitivity of WRN-resistant cell lines to irinotecan, the proteasome inhibitor MG-132, or the ATR inhibitor AZD6738 was largely unchanged, confirming WRN-specific effects.

To independently validate on-target drug resistance, we used an organoid derived from a patient with sporadic MSI colorectal cancer (CRC-14a)(6). The patient-derived organoid (PDO) was treated with HRO761 *in vitro*, and resistance emerged (Fig. 4F). WGS of the resistant PDO revealed a splice mutation (1577-2 A>T), affecting the same codon and inducing the same splice alteration observed in cell lines (Supplementary Figure 6G-H). Remarkably, HRO761-resistant organoids maintained sensitivity to VVD-133214, confirming our cell line studies (Fig. 4G-H).

During the revision of this manuscript, another WRN inhibitor, GSK4418959, was disclosed at the AACR 2025 Annual Meeting (29) and has since entered Phase I clinical trials (NCT06710847), where it is currently enrolling patients (30). The discovery and characterization of this compound will be reported in detail in a forthcoming medicinal chemistry paper. Given its potentially distinct properties, we tested its ability to overcome the F730L mutation, which emerged during selection with the combination of HRO761 and VVD-133214, and confers cross-resistance to both. Notably, cross-resistance profiling revealed that F730L-mutant cells retained sensitivity to GSK4418959 (Fig. 4B).

These findings were corroborated by additional viability assays (Fig. 4I-J and Supplementary Figure 6I). Other resistance-associated variants identified displayed diverse resistance spectra: the F730S mutation conferred resistance to HRO761 but not to VVD-133214 or GSK4418959, while G1207D and A1209D mutations appeared to confer resistance to VVD-133214 only (Fig. 4B). Finally, the G729S mutation conferred partial resistance to all inhibitors tested, though to a lesser extent with VVD-133214 and GSK_WRN3 (Fig. 4B).

These findings demonstrate that specific on-target WRN resistance mutations can confer either cross-resistance or selective resistance to individual WRN inhibitors. Notably, we demonstrate that inhibitors with distinct properties, currently in Phase I trials, can overcome resistance conferred by specific WRN mutations. This supports the use of such mutations as predictive biomarkers to guide the sequential or alternative use of mechanistically diverse WRN inhibitors.

## Discussion

Multiple WRN inhibitors are in preclinical and clinical development. The first-in-human Phase 1 trial of VVD-133214 in MSI patients demonstrated an acceptable safety profile, no dose-limiting toxicities, and anti-tumor activity across diverse histologies, establishing early clinical proof-of-concept(29). However, the emergence of resistance is likely to limit WRNi long-term efficacy. Understanding resistance mechanisms is therefore critical to refine patient stratification, enable ctDNA-based monitoring(30), and guide the design of next-generation inhibitors and rational combinations. To address this, we employed a multifaceted approach to model how MSI cells adapt to WRN inhibition, providing a first glimpse of their evolutionary trajectories under selective pressure, and identified on-target WRN mutations as a dominant driver of resistance.

To delineate the evolutionary trajectories underpinning cellular adaptation, we investigated genetic and pharmacological perturbations of WRN. Whereas complete WRN knockout caused acute loss of function and yielded no suppressor hits, pharmacological inhibition at sub-lethal concentrations partially suppressed helicase activity, allowing compensatory pathways to emerge. Our pharmacogenetic screen identified bona fide modulators of WRNi sensitivity, although none fully rescued viability. For example, loss of the MRN complex (MRE11, RAD50, NBN) or ATM reduced sensitivity to WRNi, likely reflecting a general defect in DNA damage recognition (31–33). Loss of *SMARCAL1* also conferred mild resistance by suppressing DSB accumulation at (TA)n repeats, consistent with its role in replication fork stability and recently reported interaction with TA dinucleotide repeats (22,34). WRN and SMARCAL1 may thus intersect at stalled replication forks(20,21), but with opposing roles that merit further investigation.

Semi-saturation mutagenesis predicted a spectrum of resistance variants, which aligned closely with spontaneously acquired mutations in long-term culture and were validated in vivo, including in patient-derived xenografts. While PAM and sequence constraints prevented mutagenesis of all WRN residues, our base-editing results closely mirrored those of spontaneously acquired resistance variants, highlighting the complementary value of engineered and spontaneous approaches for mapping drug resistance (25). The high mutational burden of MMR-deficient cells likely accelerates the emergence of such variants, underscoring the need to assess whether similar mutations arise in patients receiving WRNi in ongoing trials.

Resistance mutations in WRN operate through diverse mechanisms. Several clustered near or distal to drug-binding sites, where conformational changes likely impaired compound binding. A non-canonical splice-site mutation truncated exon 13 while preserving WRN function. Because this event would not have been detected by cDNA overexpression, it highlights the value of endogenous mutagenesis approaches. (35). These findings parallel known resistance mechanisms in other cancers, such as MET exon 14 skipping in NSCLC, HER2 exon 16 skipping in breast and lung cancer, BRAF(V600E) splicing in melanoma, and BRCA1 splice-site mutations in ovarian and breast cancer(36–40). In addition to splice variants, mutations clustered within the helicase domain, where conformational changes are predicted to impair inhibitor binding, and within the HRDC domain, which was selectively targeted under VVD-133214 pressure.

Whole-genome sequencing showed no significant changes in TA-repeat length in resistant cells, and no recurrent alterations, including MMR mutation reversal or counterselection (9). This contrasts with PARP inhibitor resistance, where secondary BRCA1/2 alterations restore function as a key mechanism of clinical resistance (41,42).

Collectively, our data support a model in which WRN plays an essential, non-redundant role in MSI cells due to the accumulation of TA repeat expansions across the genome. Resistance is therefore not achieved through compensatory pathways because of WRNs apparently non-redundant function. The reversion of MMR deficiency was not a resistance mechanism, presumably because it would not resolve the underlying TA repeats that underpin WRN synthetic lethality. Moreover, the presence of thousands of TA repeat expansions across the genome effectively precludes their counterselection as a possible resistance mechanism. Instead, WRNi resistance is driven by recurrent on-target mutations that impair drug binding, likely facilitated by the higher mutational burden of MMR-deficient cells. This model has direct implications for WRN-targeted therapy development and warrants investigation in clinical settings, where tumor-intrinsic evolution and stromal–immune rewiring may further shape the emergence of on-target resistance mutations.

Importantly, the spectrum of resistance mutations revealed opportunities to switch between WRNi with distinct binding modes. (43). For example, I852F variants retained sensitivity to VVD-133214, while exon 13 splice variants impaired HRO761 but not VVD-133214 binding. By contrast, F730L mutations conferred cross-resistance to VVD-133214 and HRO761 yet remained sensitive to GSK4418959 (11,44). Conversely, C727R and G729D caused cross-resistance to VVD-133214 and GSK_WRN3, while HRDC mutations (G1207D, A1209D) emerged only under pressure from VVD-133214 but retained sensitivity to the two other WRNi. These findings highlight the value of developing WRNi with complementary resistance profiles as a potential strategy to enable sequential therapy. This mirrors the clinical development of next-generation EGFR, ALK, and MET inhibitors, which target drug-resistant variants of the target proteins that evolve under drug selective pressure to overcome resistance (43,45).

Multiple WRNi are now being tested clinically, including VVD-133214 monotherapy (NCT06004245), HRO761 alone or with pembrolizumab/irinotecan (NCT05838768), and GSK4418959 with or without immune checkpoint blockade (NCT06710847). Our data suggest that resistant clones remain vulnerable to orthogonal therapies, such as chemotherapy and targeted agents (6). Notably, combining WRNi with irinotecan suppressed resistance and delayed progression, consistent with published studies(11,15). Dual WRNi with distinct mechanisms, or WRNi combined with chemotherapy, ATR inhibitors (14,28), or immunotherapy, could represent promising strategies for a more durable response, pending further validation. Notably, the preponderance of on-target resistance mutations, rather than alternative resistance mechanisms, indicates that ctDNA monitoring of hotspot mutations (e.g., G729, I852, F730L) could be an efficient approach for sensitive, early detection of resistance mutations to guide adaptive treatment strategies.

In conclusion, we present evidence that on-target WRN mutations are the principal mechanism of resistance in MSI cancer cells. These findings provide biomarkers for ctDNA-based monitoring(30), preclinical evidence that resistance mutations can be targeted with orthogonal WRNi or chemotherapy, and a rationale for combinatorial strategies to prevent or overcome resistance. By prospectively addressing resistance, this work will inform the interpretation of emerging clinical trial results and may help to accelerate the development of more effective therapies.

## Methods

### Cell models

The cell lines used in this study were sourced from the Genomics of Drug Sensitivity 1000 cell line collection and are listed in the Cell Model Passports database (https://cellmodelpassports.sanger.ac.uk/) (46). As previously reported, CRC14a patient-derived organoids were established at the Netherlands Cancer Institute and kindly provided by the lab of Emile Voest (6,47). Doxycycline-inducible WRN sgRNA-expressing cells (di-WRN) were established as reported previously. Briefly, we cloned WRN sgRNA 4 (1) into the pRSGT16H-U6Tet-(sg)-CMV-TetRep-TagRFP-2A-Hygro vector (Cellecta) and transduced Cas9-expressing HCT116 and RKO, followed by selection with 500 μg/ml hygromycin (Thermo Fisher Scientific). Single-cell clones were generated via serial dilution to ensure uniform Cas9 expression and inducible WRN-targeting sgRNA. For growth rate measurement after conditional WRN knockout induction, cells were grown in flasks with or without 2 μg/ml doxycycline for 24 hours, then seeded in 96-well plates with the same doxycycline conditions. Cell growth was monitored every 6 hours using an automated IncuCyte FLR 4X phase-contrast microscope (Essen Instruments), and the average object-summed intensity was calculated using IncuCyte software. Cas9 activity was assessed as described previously (48).

All cell lines were maintained under their original culture conditions according to supplier guidelines, with supplementation of 10% FBS, 2 mM L-glutamine, and antibiotics (100 U/mL penicillin and 100 mg/mL streptomycin) at 37 °C in a 5% CO2 incubator. Mycoplasma testing was conducted using complementary methods (MycoAlert, Lonza; EZ-PCR, Biological Industries), confirming that all cell models were free from Mycoplasma. Cell line authentication was performed using a panel of 94 single-nucleotide polymorphisms (SNPs) on a Fluidigm 96.96 Dynamic Array IFC, requiring a minimum of 75% match to the reference profile for positive authentication. Short tandem repeat (STR) profiles were also compared with those from cell line repositories. Cell models were kept in culture for an average of 36 days (up to 60 days), and all experiments were conducted within this timeframe.

### Compounds, drug screening and dose-response curve fitting

AZD66738, AZD0156, MG132, and Irinotecan were purchased from Selleck Chemicals. WRni were provided by GSK. DMSO-solubilized compounds were stored at room temperature in low-humidity (<12% relative humidity) and low-oxygen (<2.5%) environments using storage pods (Roylan Developments). Screening of cancer cell lines and organoids available at the Sanger Institute was performed using single treatments. Compounds were screened at twelve concentrations spanning a 2,048-fold range with a 2-fold dilution series. Cells were transferred into 384-well assay plates in 40 μl (cell lines) using Multidrop Combi (Thermo Fisher Scientific) dispensers. Each model’s seeding density was optimised before screening to ensure that each cell line was in the exponential growth phase at the end of the assay. Six cell densities were tested using a two-fold dilution step; each density was dispensed into 48 wells of a single 384-well assay plate and incubated for 96 hours. Cell number was quantified using CellTiter-Glo 2.0 (Promega). The maximum density tested was 3200 cells per well. Assay plates were incubated at 37 °C in a humidified atmosphere at 5% CO2 for 24 hours and then dosed with the test compounds using an Echo555 (Beckman Coulter). The final DMSO concentration was typically 0.1%. The assay plates were incubated after compound dosing, and the drug treatment duration was 72 h. CellTiter-Glo 2.0 (Promega) was added, 13.5 μl for cell lines or 20 μl for organoids, to measure cell viability. Each assay plate was incubated at room temperature for 10 min before quantification of luminescence using a Paradigm (Molecular Devices) plate reader. To estimate cell growth throughout drug treatment, an additional undrugged control plate was generated, and cell viability was measured at the time of drug treatment. These plates are referred to as a ‘day = 1’ and were repeated each time a cell line was screened. All screening plates contained negative control wells (untreated wells, n = 6; DMSO-treated wells, n = 62) and positive control wells (medium-only wells, n = 12; Staurosporine-treated wells, n = 8; and MG-132-treated wells, n = 8) distributed across the plates. These control wells were used to evaluate defined quality control criteria, including the coefficient of variation (CV) and Z-factor, as previously described (49). A maximum threshold of 0.23 was applied to the coefficient of variation (CV), and Z-factors were required to exceed a minimum threshold of 0.3. Where a cell line was sensitive to both positive controls, it had to pass Z-factor thresholds for both positive controls. Plates that did not meet these requirements were excluded from the study.

Luminescence readings were converted to cell viabilities by normalizing with reference to the DMSO-treated wells and the positive controls (viabilities of 1 and 0, respectively). Dose-response curves were then fitted to the drug-treated wells using a non-linear mixed effect model (50) to obtain IC50 estimates. Curves with a root mean squared error of greater than 0.3 were excluded from further analysis.

### CRISPR/KO genetic screens and genetic KO experiments

CRISPR screen in di-WRN cell lines (HCT116, RKO) and in SW48 treated with GSK_WRN3 was performed similarly to what was previously reported (1,51). A total of 3.3 × 10^7 of di-WRN cells were transduced with an optimized volume of the lentiviral-packaged whole-genome sgRNA library (Human CRISPR Library v1.1(1)) to achieve 30% transduction efficiency, ensuring 100× library coverage. This genome-wide library targets ∼18,000 protein-coding genes using four sgRNAs per gene, plus 1000 non-targeting controls, and was previously validated for robust dropout screening performance in cancer cell lines. The required volume was determined for each cell line by titrating the packaged library and assessing blue fluorescent protein (BFP) positivity via flow cytometry. Transductions and screens were performed in duplicates for di-WRN HCT116 and SW48, and in a single replicate for RKO cells. After puromycin selection, transduced cells were treated continuously with doxycycline (2 μg/ml) throughout the experiment. DNA was extracted for sequencing once the cells had recovered from the effects of the WRN knockout induced by doxycycline, approximately 20–30 days post-transduction, depending on the cell line. For pharmacogenomics screens, SW48 cells (3 × 10^7) were transduced in biological duplicates with the WG minimal library (51)(Gonçalves et al. 2019)(Addgene catalogue #164896) at 30% efficiency on day 1. After puromycin selection, cells were treated with either GSK_WRN3 (600 nM) or DMSO on day 10. This concentration was chosen for its ability to kill over 50% of cancer cells. At least 1.5 × 10^7 cells were maintained throughout the screening (10 days). On day 20, DNA extraction, sgRNA amplification, and sequencing were performed using 2 × 10^7 cells as the starting material. Genomic DNA was extracted using the Qiagen Blood & Cell Culture DNA Maxi Kit, 13362, as per the manufacturer’s instructions. PCR amplification, Illumina sequencing (19-bp single-end sequencing with custom primers on the HiSeq2000 v.4 platform). Experiments with an alternative WRN-targeting sgRNA described in Supplementary Figure 1 were conducted by cloning a different sgRNA sequence (GAGCATGAGTCTATCAGAT) into the lenti-sgRNA neo plasmid (Plasmid #104992). Cells were transduced and selected accordingly. To genetically inactivate TP53, SMARCAL1, SUPT7L, MUS81, and TADA1, di-WRN- or Cas9-expressing cell lines were transduced overnight with lentiviral constructs containing specific sgRNAs, in the presence of polybrene (8 μg/mL). The sgRNAs used were as follows: SMARCAL1 (SW48: GCGCTGTCGAGCAGCTATGC, HCT116: GGCAGCTATGCCGGTCCTAA), MUS81 (HCT116: GGTGCTGTATCGATCCACCA, SW48: GAGCGGCACCGAACATCGGG), SUPT7L (GCACAGTCAAAGCCCGCGT for both cell lines), and TADA1 (ACAACGCGTGAGAATGGCC for both cell lines). Medium was refreshed for a fresh complete medium the following day. Cells were treated with blasticidin (20 μg/mL) and puromycin (2 μg/mL, Thermo Fisher Scientific, A1113803), hygromycin 500ug/mL (Invitrogen; Cat#10687010) or Geneticin (Invitrogen; Cat# 10131-035), depending on the respective selective agent, to select for Cas9-expressing cells carrying the sgRNAs.

### CRISPR–Cas9 Base Editing Screens

Base editing experiments were performed as recently reported, using the same Base editing library that targets the WRN gene (10). Briefly, we introduced base editing machinery into KM12 and RL-95-2 cells through co-transfection with FuGENE HD (Promega), utilizing a plasmid encoding Cas9 and a gRNA targeting the CLYBL locus (5′-ATGTTGGAAGGATGAGGAAA-3′). This was paired with a plasmid containing a tet-ON base editor, blasticidin resistance, and mApple expression cassettes within CLYBL homology arms, following methods reported previously. To boost homologous recombination (HR) rates, cells were pre-treated overnight with 1 μM DNA-PK inhibitor AZD7648. Post-transfection, cells were selected with 10 μg/mL blasticidin (Thermo Fisher Scientific) for four days, then maintained with 5 μg/mL. Cell pools were further refined via FACS for mApple expression. Base editing efficiency was assessed using BE-FLARE (52). Clonal lines of BE3.9max NGN and ABE8e NGN were employed for detailed screening. We conducted base editing screens using the same sgRNA library recently reported (10), sgRNA representation of 1,000-fold, utilizing viral doses that resulted in 30% to 50% cell infection. Postinfection, cells were selected with puromycin for 4 days. Five days postinfection, a baseline (T0) pellet was collected. Base editing was induced by administering doxycycline (1 μg/mL) for 3 days. Following a 10-day selection phase aimed at removing sgRNAs essential for cell growth, the cells were maintained in culture at a minimum density of 15 million cells per three-layer flask. They were split regularly to support optimal growth. After this phase, cells were divided into control and drug-treated arms. KM12 cells in the drug-treated arm received 850 nM of GSK_WRN3 and 120 nM of HRO761, while RL95 cells were treated with 350 nM of GSK_WRN3. CBE and ABE treatments were administered at the following concentrations for each cell line. This setup enabled us to assess the differential responses in the control and drug-treated arms over an additional 10-day period. Each screen was independently replicated 2 times on separate days for consistency. Cell pellets were processed to extract DNA for DNA analysis, and the gRNA cassette was amplified through PCR. This was done in parallel to maintain library diversity. After post–column purification, the PCR products were indexed and sequenced on the Illumina HiSeq 2500 using 19 bp single-end sequencing and a custom primer. We employed BEstimate to analyze screen results and annotate gRNAs. We assumed a wild-type genome as the baseline, disregarding specific SNPs or SNVs in the two cancer cell lines. We normalized read counts to reads per million, adding a pseudo-count for calculation purposes. We averaged reads per million values from replicates for consistent gRNA analysis. Average log2fold-change values were calculated, and z-scores were derived by normalizing them against their SD. sgRNAs with fewer than 100 reads in the initial samples were also excluded from downstream analysis.

### Generation of isogenic SW48 cell lines

Isogenic SW48 colorectal cancer cell lines carrying defined WRN SNPs were generated using CRISPR–Cas9 editing. Cells were nucleofected using the Amaxa 4D Nucleofector System with the Lonza SF kit (PE kit, program DP100). For cell line editing, RNP complexes contained Cas9 protein, dual synthetic sgRNAs targeting WRN exon 13 or exon 21, and single-stranded oligonucleotide donor template (ssODNs) encoding the desired nucleotide substitutions. Transfection efficiencies were monitored using a GFP control plasmid electroplated separately and analysed through flow cytometry. Edited bulk populations were assessed for HDR efficiency by T7/TIDER assays, and upon successful editing, they were sorted for single-cell clones into 96-well plates through FACS sorting. Clones were expanded, genotyped by PCR, and validated by Sanger sequencing to confirm the presence of heterozygous or homozygous knock-in alleles. For both WRN mutations, heterozygous clones were employed to generate the drug sensitivity data presented in the heatmap.’

### Cell viability assays

Drug sensitivity assays were conducted to confirm the resistance profiles of each cell line. Approximately 1.5–2.5 × 10³ cells per well were seeded into 96-well plates. The following day, a range of drug concentrations was added, with each concentration tested in triplicate for each line. Cells were incubated for 7–10 days, after which viability was assessed using the CellTiter-Glo 2.0 Assay (Promega, G9241). For viability measurement, 25 μL of CellTiter-Glo 2.0 reagent was added to each well, followed by a 20-minute incubation at room temperature in the dark. Luminescence was then measured using an Envision Multiplate Reader.

### Clonogenic assays

For clonogenic growth assays, 2.5-6 × 10³ cells per well (depending on the cell model) were seeded into 6-well plates and treated with different drug concentrations. Colonies were fixed with 3% paraformaldehyde and 1% glucose, and stained with 0.5% Crystal Violet solution (Merck, Cat. no. C0775) to visualize colony formation.. Colony formation was assessed 10–14 days after treatment.

### TTP Assay

For time-to-progression assays, 1 × 10⁷ cells were plated in their respective growth media with WRN inhibitors GSK_WRN3 or HRO761. At each time point, cultures were harvested, viable cells were counted, and all available viable cells were reseeded into fresh medium containing the same drug concentration. Counts of zero indicate time points when the number of viable cells was too low to allow reseeding; in such cases, only medium and drug were refreshed.

### RNA Interference–Based Sensitivity Assay

Experiments involved seeding approximately 1.5–3.5 × 10³ cells per well in a 96-well plate, followed by reverse transfection with ON-TARGETplus siRNA at a final concentration of 20 nmol/L, using RNAiMAX (Invitrogen) according to the manufacturer’s protocol. Each experiment included controls: transfection reagent only (mock control), a nontargeting pool (negative control, Dharmacon, D-001810–10–05), a polo-like kinase 1 (PLK1) pool (positive control, Dharmacon, L-003290–00–0010), and a WRN-targeting pool (Dharmacon, L-010378–00–0005). The siRNA sequences were: nontargeting (UGGUUUACAUGUCGACUAA, UGGUUUACAUGUUGUGUGA, UGGUUUACAUGUUUUCUGA, UGGUUUACAUGUUUUCCUA), PLK1 (GCACAUACCGCCUGAGUCU, CCACCAAGGUUUUCGAUUG, GCUCUUCAAUGACUCAACA, UCUCAAGGCCUCCUAAUAG), and WRN (GAUCCAUUGUGUAUAGUUA, GCACCAAAGAGCAUUGUUA, AUACGUAACUCCAGAAUAC, GAGGGUUUCUAUCUUACUA). Cells were cultured for 5–7 days, and viability was measured using the CellTiter-Glo 2.0 Assay (Promega, G9241).

### Western blot

Western blotting confirmed WRN depletion and assessed DNA damage in cells treated with varying concentrations of multiple WRN inhibitors in different drug-resistant models. For the blots, 4-8 × 10^6 cells were cultured in 10 cm dishes and treated with GSK_WRN3, HRO761, VVD-133214 or DMSO. Cells were lysed 48 hours post-treatment using 100-150 µL of RIPA buffer containing protease inhibitors. Lysate concentrations were determined using the BCA Assay. Each sample, containing 20-30 µg of lysate, was run on a 4-12% Bis-Tris gel (Invitrogen) and transferred to a PVDF membrane. Membranes were blocked with 5% milk in TBST and incubated overnight with primary antibodies: anti-WRN (ABcam, AB124673, 1:1000), anti-P-Histone H2AX (NB100-384, 1:1000), and anti-β-tubulin (Sigma-Aldrich, T4026, 1:5000) as a loading control. MUS81 was detected using the Abcam antibody (Cat. ab14387) at a dilution of 1:1000. TADA1L was detected with the Invitrogen antibody (Cat. PA5-69843) at a dilution of 1:1000, and SUPT7L was identified using the Invitrogen antibody (Cat. PA5-70808) at a dilution of 1:1000. Blots were washed and incubated with an anti-Rabbit IgG HRP-linked secondary antibody (GE Healthcare, #NA931V-ECL HPR) for 1 hour at room temperature. After a final wash in TBST, signals were detected using Super Signal Dura. Precision All Blue Plus Protein Standards (BioRad, cat. 1610373) were used as molecular weight markers.

### WRN inducible knockout in HCT116 mouse xenograft tumors

In vivo experiments with inducible di-WRN HCT116 were performed as previously reported (1). Female NOD/SCID mice (Charles River Laboratories) were used for all in vivo studies. All procedures were approved by the Ethics Committee and the Italian Ministry of Health (authorisation 806/2016-PR), adhering to the agreed-upon guidelines. Mice were maintained in hyperventilated, pathogen-free conditions in individually sterilised cages with up to seven mice per cage and provided with sterilised food, water, and bedding. Inducible WRN sgRNA-expressing HCT116 cells xenografts were established by subcutaneously injecting 2 × 10^6 cells into the right posterior flank of 5- to 6-week-old mice. Tumor size was measured with callipers, and volume was calculated using the formula 4/3π × (d/2)^2 × (D/2), where d is the minor axis, and D is the major axis. When tumors averaged 250–300 mm^3, animals were randomized by tumor size. Doxycycline (Sigma-Aldrich, D9891) was administered orally by gavage at a dose of 50 mg/kg daily. Each experimental group consisted of 8–10 mice to estimate within-group variability. Allocation to treatment groups was done during randomization, with measurements taken blind. The maximum tumor volume allowed was 3,500 mm^3, which was not exceeded. In vivo procedures and data were managed using the Laboratory Assistant Suite for automated data tracking and analysis.

### In Vivo Studies in SW48 Xenografts and Patient-Derived Xenografts (PDX)

All procedures related to animal handling, care, and treatment in this study adhered to guidelines approved by the Institutional Animal Care and Use Committee (IACUC) and followed the standards of the Association for Assessment and Accreditation of Laboratory Animal Care (AAALAC). The in vivo study was conducted in accordance with GSK’s 3Rs principles (Replacement, Reduction, Refinement). Experimental protocols for SW48 were approved by the GSK IACUC and performed at the GSK US Cambridge site. For the tumor xenografting process, 5 × 10^6 SW48 cells were mixed in a 1:1 ratio with RPMI and Matrigel (100 µL) and xenografted into the right flank of 6- to 8-week-old female Crl nu mice (Strain #088, Charles River Laboratory, Wilmington, MA, USA), weighing 20–25g. PDS-0023 PDX samples, sized 3mm x 3mm, were implanted using a 10G Trocar into the right hind flank of NOD.Cg-prkdcscid IL2rgtmWjl/Szj mice (Strain # 005557, Jackson Lab, Bar Harbor, ME USA), aged 6-8 weeks and weighing over 17g. Once tumors reached approximately 80–160 mm³, animals were randomly assigned to groups based on individual tumor size using a stratified randomization method for SW48. Body weight and tumor volume were measured twice weekly using an electric calliper (Fowler Ultra Cal V) and an Ohaus electronic scale (STX421), with data automatically recorded in the Study Log (4.6). Tumor volume was calculated as 0.5 x L x W^2, where L represents tumor length (longest dimension) and W represents width (perpendicular to L). HRO761 was formulated in 0.5% methylcellulose (400 cp) with 0.5% Tween 80 and administered via oral gavage at 60 or 120 mg/kg in a 10 mL/kg volume once daily, while the vehicle control group received 10 mL/kg of 0.5% methylcellulose (400 cp) with 0.5% Tween 80 for 100 days. Tumor samples were collected and snap-frozen 24 hours post-final dose at the end of the study.

DNA was extracted. Explants were sequenced using the Novogene Precision Medicine 2.0 (NovoPM 2.0) gene panel. Following extraction, DNA quantity was measured with the Qubit® (Life Technologies, Thermo Fisher Scientific, USA). Agarose gel electrophoresis (Bio-Rad Laboratories, USA) was then used to assess any contamination or degradation of the DNA samples. Library preparation for NovoPM™ 2.0 was conducted using custom IDT probes and Novogene’s in-house reagents. Standard protocols were followed, with library quantification assessed via qPCR and quality confirmed using NGS3K (PerkinElmer, Inc., USA). Sequencing was performed on the Illumina NovaSeq 6000 platform (Illumina, USA) using a paired-end (PE) 150 sequencing strategy, following standard user manual procedures. Quality control of sequencing data was conducted using Novogene’s in-house pipeline, which generated clean data for bioinformatics analysis. The Burrows-Wheeler Aligner (BWA) was used to align paired-end clean reads to the human reference genome (hg19). Samtools was employed for sorting, and duplicates were marked with Picard, with the alignment results stored in BAM format. Coverage and depth calculations were based on the final BAM files. Somatic mutations were analyzed using Novogene’s in-house algorithm, NovogeneSGZ, which predicts the origin of short variant mutations (germline or somatic) using data solely from tumor samples.

### Immunohistochemistry

Formalin-fixed, paraffin-embedded tissues from xenografts were sectioned (10-μm thick), with 4-μm sections dried overnight at 37 °C. Slides were deparaffinized in xylene and rehydrated through a graded series of alcohols to water. Endogenous peroxidase was blocked with 3% hydrogen peroxide for 30 min. Antigen retrieval was performed in 10 mmol/l citrate buffer (pH 6.0) using a microwave (750 W for 10 min). Slides were incubated with monoclonal mouse anti-human KI-67 (1:100; DAKO) or polyclonal WRN antibody NBP1-31895(1:100, Novus) overnight at 4 °C. After washing in TBS, slides were treated with anti-mouse secondary antibody (DAKO Envision+System HRP-labelled polymer) for 1 h at room temperature. Immunoreactivity was detected using DAB chromogen (DakoCytomation Liquid DAB Substrate Chromogen System, DAKO) for 10 min. Slides were counterstained with Mayer’s haematoxylin, dehydrated, cleared in xylene, and mounted with DPX (Sigma-Aldrich). Negative controls omitted the primary antibody. Stained slides were scanned at 40× magnification. For Ki-67, ten images from three cases were analyzed with ImageJ (NIH) to segment cells with positive and negative nuclei. The percentage of positively stained cells was calculated, verified by visual inspection for accuracy.

### TrAEL-seq for DSB mapping

For TrAEL-seq, 2-3 million cells were seeded in T150 flasks. Two days later, these cells were treated with GSK_WRN3 (2 uM) or HRO761 (0.5 uM) for 24h. Cells were harvested and counted 24 hours post-treatment. 1-2 x 10⁶ cells per treatment were washed with PBS, centrifuged, and resuspended in 1mL of cold L Buffer (100mM EDTA pH8, 10mM Tris pH 7.5, 20mM NaCl) before embedding in agarose for DNA extraction. DNA ends were tailed with ATP and ligated to adaptors carrying in-line indexes, then pooled and processed into a single TrAEL-seq library as described(24). Libraries were sequenced on an Illumina NextSeq 500 as High Output 75 bp Single End by the Babraham Institute Next Generation Sequencing facility, then reads were trimmed, deduplicated, de-multiplexed into individual libraries, and mapped to GRCh38 by the Babraham Institute Bioinformatics Facility using scripts available at https://github.com/FelixKrueger/TrAEL-seq.

### Analysis of TrAEL-seq data

Mapped reads were imported into SeqMonk v1.48 (https://www.bioinformatics.babraham.ac.uk/projects/seqmonk/) and truncated to 1 nucleotide at the 5′ end, representing the last nucleotide 5′ of the strand break. For an unbiased set of cleaved TAs in SW48 cells, all TA tracts (3) to which 3+ reads mapped in each drug were used (compared to a genome-wide background of 0-1 read for features of this size), 93% of peaks called for GSK_WRN3 were also called for HRO761. Normalisation coefficients were calculated based on the set of TA tracts not called as peaks. For the graph of TA-rich peaks in SW48, read counts were calculated for all TA tracts, normalised then log2 transformed and signals quantified across the set of cleaved TAs for the relevant drug. For metaplots, read counts were performed across TAs and in 10bp windows at 1bp spacings over 100bp either side, normalised as above and averaged to generate metaplots. Equivalent metaplots generated on the non-cleaved TA set showed no enrichment in any condition. Final graphs were plotted using GraphPad Prism 10.1.1.

### DNA and RNA Sequencing

DNA and RNA were extracted from cancer cell lines and organoids using the Qiagen AllPrep DNA/RNA Mini Kit. Whole-genome sequencing libraries were prepared, and paired-end reads (150 bp) were generated on the Illumina HiSeq X Ten platform, achieving an average sequencing depth of 30x.

### Data analysis

For WG CRISPR-Cas9 screenings, sgRNAs with zero read counts in control samples were excluded. Log2 fold changes (L2FC) were calculated from normalized read counts, achieved by dividing sgRNA reads by total sample reads, multiplying by 1,000,000, and adding a pseudocount of 1 to prevent division by zero. We determined the average fold change for each gene by taking the mean of the fold changes across all sgRNAs targeting that gene. To combine replicate data, we averaged these gene-level fold changes. For the MAGeCK analysis, we used default settings with one modification: the normalization parameter was set to ‘none,’ as normalization had already been applied to the corrected input counts. To annotate hits from the pharmacogenomic screen, genes conferring resistance were selected based on fold change (FC) > 1.5 and –log₁₀(p-value) > 4. The resulting gene list was analyzed for functional enrichment using the g:Profiler tool(53) (https://biit.cs.ut.ee/gprofiler) with default parameters. Enrichment was assessed across Gene Ontology categories, including Biological Process (BP), Molecular Function (MF), and Cellular Component (CC). Significant GO terms (adjusted p-value < 0.05) were visualized in a heatmap to highlight key pathways associated with WRN inhibitor resistance.

For WGS, whole-genome paired-end sequencing reads (150 bp) were generated using the Illumina HiSeq X Ten platform. Reads were aligned using the Burrows-Wheeler Alignment (BWA-MEM) tool(54). PCR duplicates, as well as unmapped and non-uniquely mapped reads, were filtered out before proceeding with downstream analysis. Single-base substitutions and indels were identified using CaVEMan (55) and cgpPindel (56). Common germline variants were filtered out using in-silico normal sample generated using GRCh38 ref genome, additional germline variants and technology-specific artefacts were removed by filtering against the panel of 100 unrelated normal samples (ftp://ftp://ftp.sanger.ac.uk/pub/cancer/dockstore/human/GRCh38_hla_decoy_ebv/SNV_INDEL_ref_GRCh38_hla_decoy_ebv-fragment.tar.gz) and gnomAD-v3.1.1 (57) and 1000 Genome variants (58) with AF>0.001. Additional post-processing filters were applied using the in-house post-processing tool cgpCaVEManPostProcessing (https://github.com/cancerit), and variant sites flagged as ‘PASS’ were considered for further downstream analysis. Unbiased analysis of mutant and wild-type reads found at the loci of the base substitutions and indels were assessed within a group of samples for a given Cell line using vafCorrect (59). For the purposes of defining resistance-associated events, only WRN mutations with a variant allele frequency (VAF) >0.03 were included in the analysis. The Jaccard index (Intersection over Union), which measures the similarity between two samples, was calculated for all the PASS variants within a pair of related samples. Tumor mutation burden is assessed using the number of germline mutations found per Mb of effective genome size. For all WGS data, the fragments per base per million were calculated in each interval using the method previously reported (3). Median values for broken and unbroken regions were then estimated for each sample. TA repeat expansion was assessed by measuring the normalized coverage depth at a predefined set of reference broken and non-broken TA repeat regions (3). Paired-end transcriptome reads (75 bp) were quality filtered and aligned to the GRCh38 reference genome (Ensembl build 98) using STAR v2.5.0c(60) with default parameters (https://github.com/cancerit/cgpRna). The resulting BAM files and splice junctions were visualized in JBrowse(61). The plots and graphs were generated using GraphPad and Spotfire software.

### Molecular Modeling and Structural Analysis of WRN Mutations

Structures of WRN bound to both the reversible inhibitor HRO761 (pdb 8PFO) and covalent inhibitor VVD-133214 (pdb 7GQU) were downloaded from the RSCB PDB database and prepared using Maestro with default parameters, Schrodinger Suite v2024.02. Mutations I852F and G729D were manually modelled in crystal structures of the two known binding modes. Local energy minimization for the area surrounding the mutation was performed using Prime tools (62) in Schrodinger Suite v2024.02. Binding pocket surface calculation (4.5Å cutoff from ligand) and molecular visualization was done in MOE v2022.02 (63).

## Data availability

TrAEL-seq sequencing data have been deposited in the Gene Expression Omnibus (GEO) repository under the accession number GSE279461.

## Acknowledgements

We thank the Garnett laboratory, the Gene Editing team, and the drug screening teams at the Sanger Institute for their assistance and data generation. The authors wish to acknowledge the contribution of the CASM Support team at the Wellcome Sanger Institute. We thank Emile Voest for providing organoid models and for helpful discussion of the results. This research was funded in whole, or in part, by Wellcome Trust Grant 206194. Funding was also provided by Fondazione AIRC per la Ricerca sul Cancro, AIRC 5×1000 grant 21091 (to A.B. and L.T.). M.A.C. is supported by Cancer Research UK [RCCCDF-Nov23/100002]. E.G. work was supported by national funds through FCT, Fundação para a Ciência e a Tecnologia, under project UIDB/50021/2020 (DOI:10.54499/UIDB/50021/2020). E.G. work is supported by national funds through Fundação para a Ciência e a Tecnologia, I.P. (FCT) under projects UID/50021/2025, UID/PRR/50021/2025, SARC-RON-AI, through RE-C05-i08.M04), and SYNTHESIS. JH and KM were supported by BBSRC BI Epigenetics ISP BBS/E/B/000C0523 and BB/W509917/1, Babraham Institute’s BBSRC Core Capability Grant (CCG) BB/CCG2310/1 and Institute Development Grant BB/IDG2310/1. GlaxoSmithKline have awarded M.J.G. and G.P. research grants. TrAEL-seq libraries were sequenced by Babraham Genomics Facility - Geno06 and data processed at Babraham Bioinformatics Facility – Bioinf01. Figure components were created with BioRender.com.

## Author contributions

GP, YR, BS, and MJG conceptualized the study. GP, AAS, SV, SW, RB conducted experiments. LT and AB supervised the in vivo CRISPR experiments, while FS conducted the IHC analyses. Drug screening was performed by SB, EM, TB, ZW, HGD, AK, MGC, and FG. HL performed curve fitting. Data analysis was conducted by GP, SB, MC, and EK. GS, YL, EB, ND, and RS designed and performed in vivo experiments. GVG and IBH contributed to molecular modeling and structural analysis of WRN mutations. MC provided key reagents, expertise for base editing experiments, and assisted with data analysis. MS cloned the base editing sgRNA library. The gRNA library design was a collaborative effort by KMcC, MC, and GP. EV contributed key reagents. JH and KM performed and analyzed TrAEL-seq experiments. MD and PL assisted with the synthesis of the compound. All IDEAYA Biosciences authors (BJ, DM, JP, PB, JT, MW) contributed to compound development, screening, or data interpretation. The manuscript was written by GP and MJG, with review and input from all contributing authors.

