## Supplementary Material for "On-target mutations confer resistance to WRN helicase inhibitors in Microsatellite Unstable Cancer Cells"

**Supplementary Figure 1**

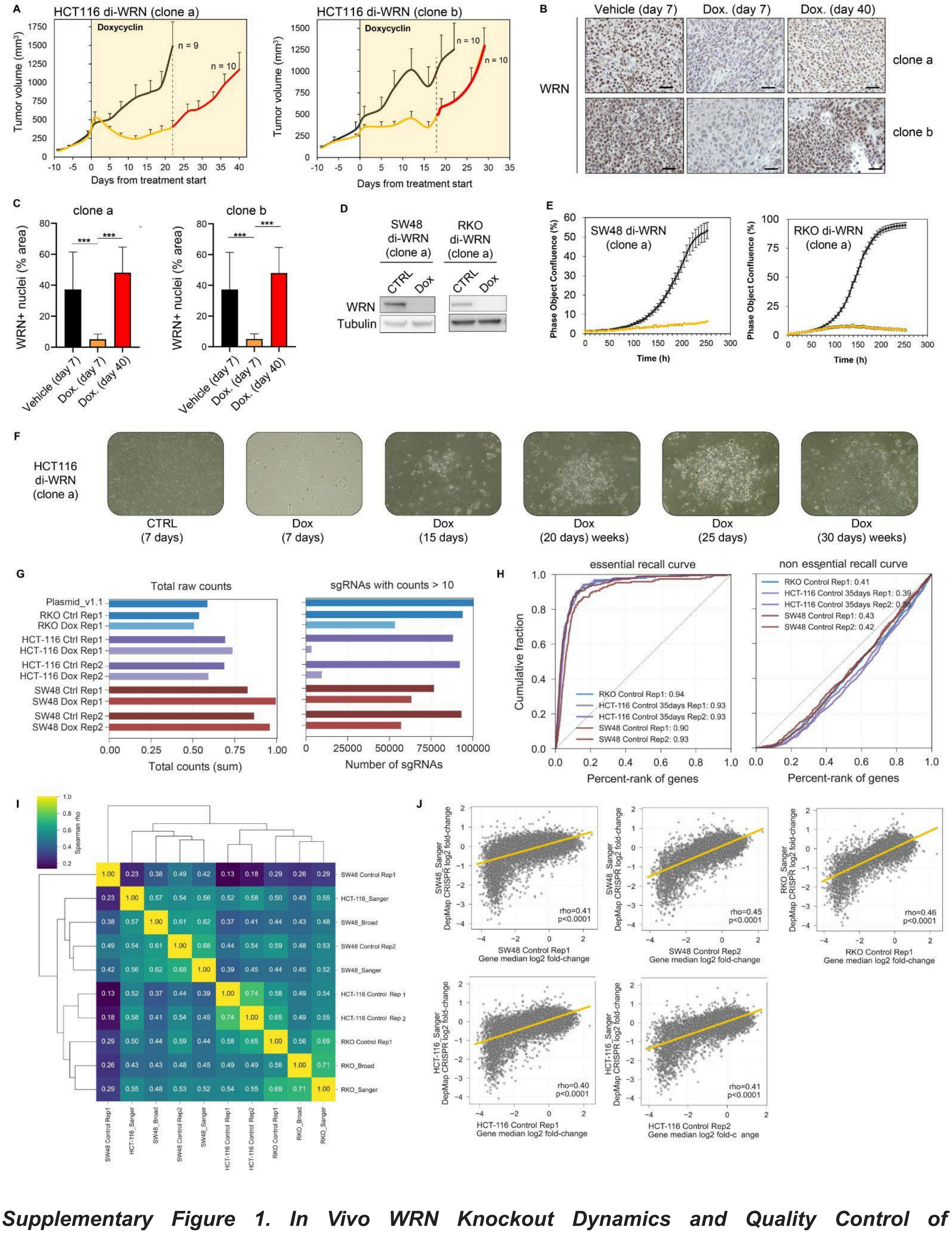


***Supplementary Figure 1. In Vivo WRN Knockout Dynamics and Quality Control of Genome-wide CRISPR Screens in MSI Models with Inducible WRN Knockout.*** *(A) Tumour volume of HCT116 di-WRN xenografts (clone a, left; clone b, right) treated with doxycycline (yellow) or vehicle (grey) for 21 days, as published in doi.org/10.1038/s41586-019-1103-9. Extended monitoring up to day 40 (red) is unpublished data. Data are mean ± s.e.m.; cohort sizes are indicated in each graph. (B) WRN immunohistochemistry (IHC) of HCT116 di-WRN tumours (clone a: top row; clone b:*

*bottom row) collected at days 7 (early) and 40 (relapse) following doxycycline or vehicle treatment. Scale bar: 50 µm; magnification 40x. (C) Quantification of WRN IHC signal from the panel B. Data are mean ± s.d. of 10 fields from three independent tumours (n = 30). P-values were calculated by two-sided Welch's t-test (***p < 0.001). (D) Western blot analysis of WRN protein levels following doxycycline-induced knockout in SW48 and RKO di-WRN subclones. Tubulin was used as a loading control. Representative of two independent experiments. (E) Growth rate curves of SW48 and RKO di-WRN subclones in the absence (black) or presence (yellow) of doxycycline (2 µg/ml). Mean ± s.d. of 10 technical replicates per condition; representative of two experiments. (F) Representative images of resistant clones emerging after doxycycline administration in di-WRN lines transduced with a genome-wide CRISPR knockout library. (G) Total read counts (left) and sgRNAs with >10 counts (right) across samples from genetic screens, demonstrating library coverage and consistency. (H) Essential and non-essential gene recall curves benchmark screen performance across samples using reference gene sets. (I) Pairwise Spearman correlation heatmap of gene-level CRISPR screen effects across cell lines, replicates, and reference datasets (Project SCORE and DepMap). Correlation coefficients validate screen quality and inter-dataset reproducibility. (J) Scatter plots comparing gene-level log2 fold changes (Control vs. Plasmid) from di-WRN CRISPR screens to corresponding data from Project SCORE. Each plot displays the Spearman correlation coefficient (ρ), highlighting rank-based concordance across datasets.*

### Supplementary Figure 2

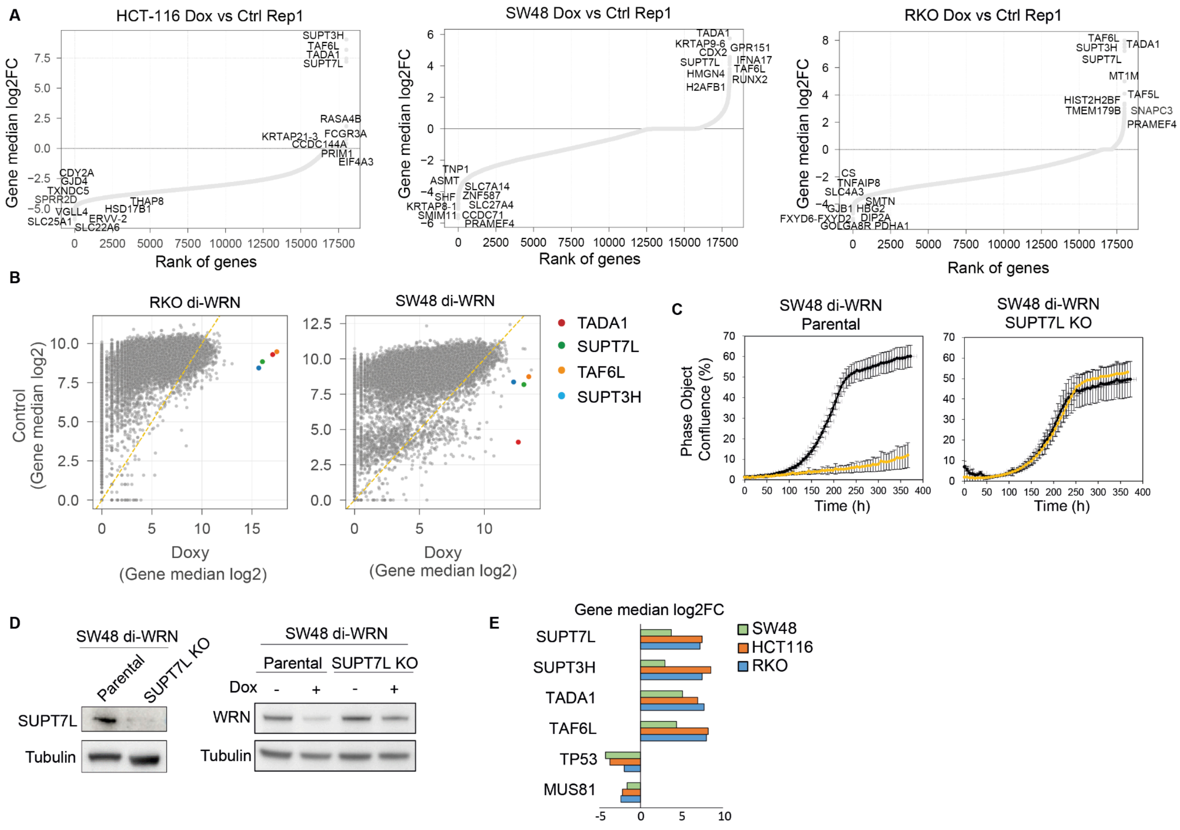


***Supplementary Figure 2. Results and Validation of Genome-wide CRISPR Screens in Inducible WRN Knockout MSI Cancer Models.*** *(A) Rank-ordered gene-level median log2 fold changes for doxycycline vs control conditions in HCT116, SW48, and RKO (bottom) cells. SAGA complex members (e.g., TADA1, SUPT3H, TAF6L, SUPT7L) are among the top-scoring hits. (B) Gene-level*

*quantification of sgRNAs in doxycycline-treated cells versus controls in RKO and SW48 di-WRN cells. Recurrent enriched resistance-associated genes are colored. (C) Growth rate curves of SW48 di-WRN parental and SUPT7L KO cells in the absence (black) or presence (yellow) of doxycycline. Mean ± s.d. of 10 technical replicates; representative of two experiments. (D) Western blot showing WRN and SUPT7L protein levels in SW48 di-WRN parental and SUPT7L KO cells ± doxycycline. Tubulin is a loading control. (E) Bar plot summarising gene-level log2 fold change (median across sgRNAs) for WRN modulators (TP53, MUS81) and SAGA complex members in SW48, HCT116, and RKO cells.*

### Supplementary Figure 3

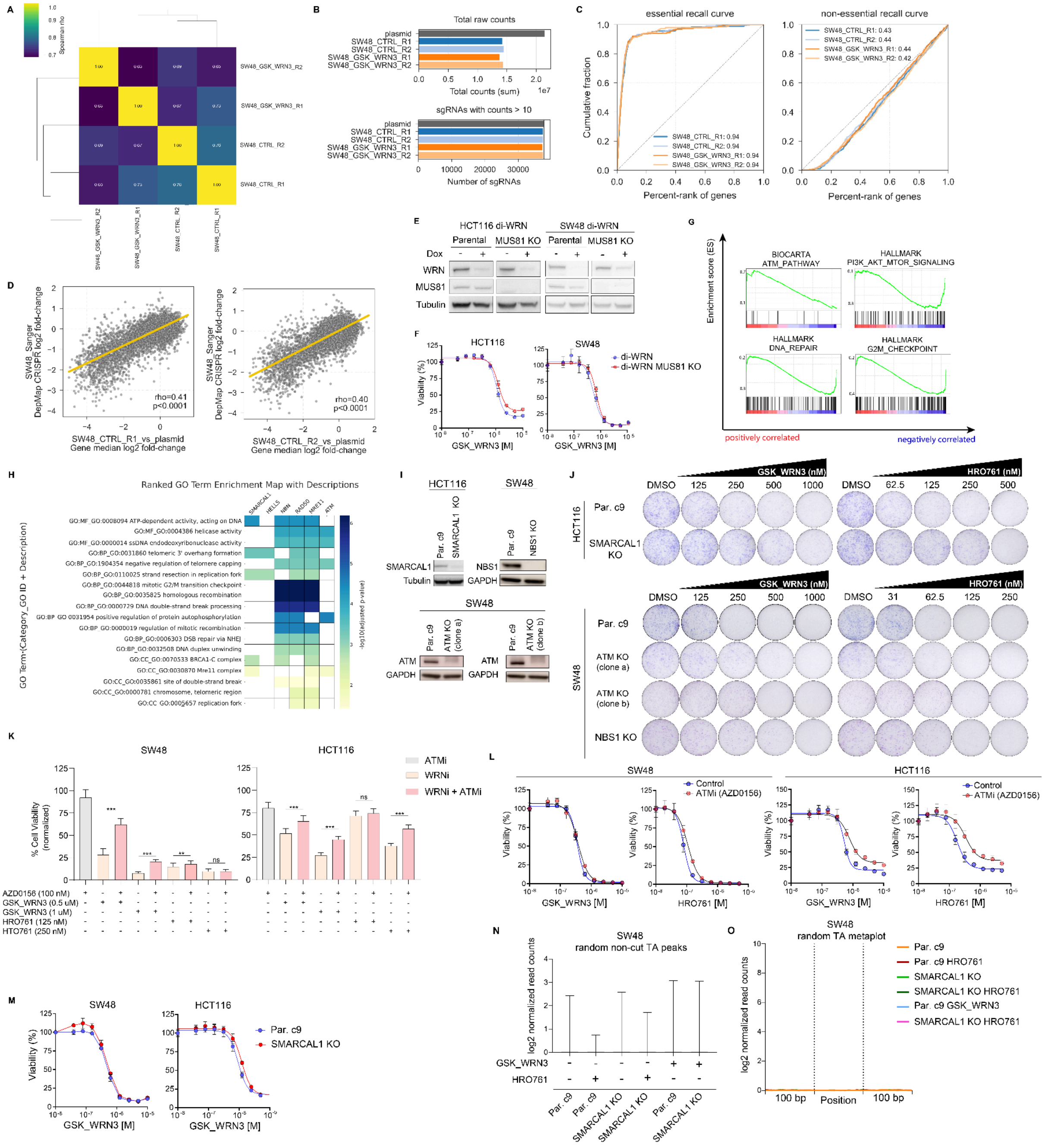


***Supplementary Figure 3. Results and Validation of Genome-wide CRISPR Screens in MSI Cancer Models Treated with WRN Inhibitors.*** *(A) Clustering of samples in the CRISPR/Cas9*

*pharmacogenomic screening in SW48 cells. The heatmap legend indicates Spearman correlation.* **(***B) Total read counts (top) and sgRNAs with >10 counts (bottom) across samples from genetic screens, demonstrating library coverage and consistency. (C) Essential and non-essential gene recall curves benchmark screen performance across samples using reference gene sets. (D) Scatter plots comparing gene-level log2 fold changes (Control vs. Plasmid) from di-WRN CRISPR screens to corresponding data from Project SCORE. Each plot displays the Spearman correlation coefficient (ρ), highlighting rank-based concordance across datasets. (E) Western blot analysis showing WRN and MUS81 protein levels in HCT116 and SW48 di-WRN cells. Cells were treated with doxycycline (Dox) to induce WRN knockdown, and MUS81 knockout (KO) was confirmed. Tubulin is shown as a loading control. (F) Dose-response curves relative to control for HCT116 and SW48 di-WRN cells with or without MUS81 knockout treated with GSK_WRN3. Data are average ± SD of three technical replicates and represent three independent experiments. (G) Gene set enrichment analysis (GSEA) of pathways differentially enriched based on log2 fold-change ranked values from the chemogenomic screen in SW48 cells treated with GSK_WRN3. (H) Genes conferring resistance to WRN inhibition were identified via a pharmacogenomic CRISPR screen and annotated using* the g:Profiler *tool. The heatmap shows significantly enriched GO terms (rows) and associated genes (columns), with color intensity representing* –$\log_{10}$ *adjusted p-values. Enrichment highlights key processes, including DNA repair, telomere maintenance, and the response to replication stress. (I) Western blot for SMARCAL1 in HCT116 and SW48 parental cells and their corresponding SMARCAL1, ATM, and NBS1 knockout (KO) derivatives. Tubulin and GAPDH are used as controls. (J) Clonogenic assays evaluating the sensitivity of HCT116 and SW48 parental cells and SMARCAL1, ATM, and NBS1 isogenic knockout clones to increasing concentrations of GSK_WRN3 (left panels) and HRO761 (right panels). DMSO-treated cells served as controls. (K) Percentage cell viability (normalised to control) for SW48 (left) and HCT116 (right) MSI cell lines treated with WRN inhibitors (GSK_WRN3 and HRO761) with or without 6-hour ATMi (AZD0156, 100 nM) pre-treatment. Error bars represent the standard deviation (SD) of six technical replicate wells per plate, from two independent biological replicates (n = 2). Statistical significance was assessed using a paired t-test (*$p < 0.05$; **$p < 0.01$; **$p < 0.001$; ns = not significant). (L) Dose-response curves for SW48 and HCT116 parental cells treated with GSK_WRN3 and HRO761, either alone or following a 6-hour pre-treatment with ATMi (AZD0156, 100 nM). The total assay duration was 8 days. Data represent the mean ± SD of three technical replicates and are representative of three independent experiments. (M) Dose–response curves of HCT116 and SW48 isogenic cell lines, with or without SMARCAL1 knockout, treated with GSK_WRN3 for 96 h. The data are plotted as a percentage of viability relative to the DMSO control. Data are average ± SD of three technical replicates and represent three independent experiments. (N) Quantification of TrAEL-seq read counts at random non-cut TA peaks in SW48 cells under various conditions, as indicated. (O) Metaplot analysis of TrAEL-seq data centred around TA dinucleotide repeats, comparing parental c9 and SMARCAL1 knockout SW48 cells treated with GSK_WRN3 or HRO761.*

### Supplementary Figure 4

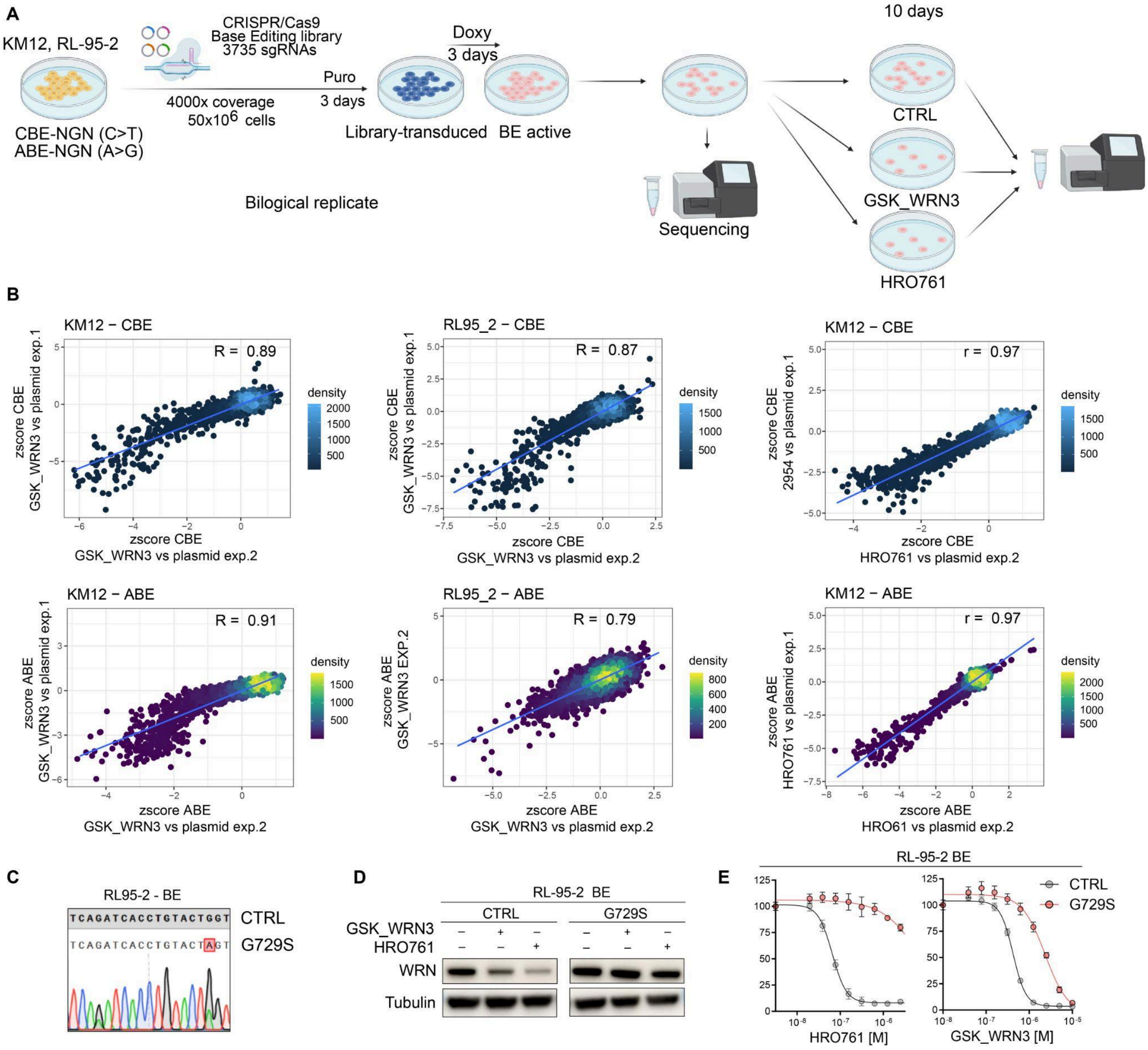


***Supplementary Figure 4. Comparative Analysis of Base Editing Screens in KM12 and RL95-2 Cells Treated with GSK_WRN3 and HRO761.*** *(A) Schematic of experimental setup: KM12 and RL95-2 cell lines were subjected to CRISPR/Cas9 base editing using a library containing 3759 sgRNAs with either CBE-NGN (C>T) or ABE-NGN (A>G) editors. Cells were transduced with the base editing library, selected with puromycin for three days, and then induced with doxycycline for three additional days to activate base editing. Cells were treated with either GSK_WRN3 (850 nM for KM12 and 350 nM for RL95-2) or HRO761 (120 nM) or left untreated as a control for ten days before sequencing. (B) Comparison of z-scores for GSK_WRN3 and HRO761 treatment across two biological replicates for CBE (top row) and ABE (bottom row) in KM12 and RL95-2 cells. High correlation values (R-values) were observed, indicating reproducibility across experiments. While KM12 cells were screened with GSK_WRN3 and HRO761, RL95-2 cells were only screened with HRO761. (C) Representative chromatograms of Sanger-sequenced PCR products confirming G729S mutations introduced via CRISPR-Cas9 base editing in KM12 and RL95-2 cells. D) Western blot analysis of RL95-2 cells comparing isogenic parental and G729S cells treated with HRO761 and GSK_WRN3. Tubulin serves as the loading control. Results are representative of two independent experiments. E) Dose-response viability curves in RL95-2 cells comparing parental and G729S mutant cells treated with HRO761 and GSK_WRN3. Data are average ± SD of three technical replicates and represent three independent experiments.*

**Supplementary Figure 5**

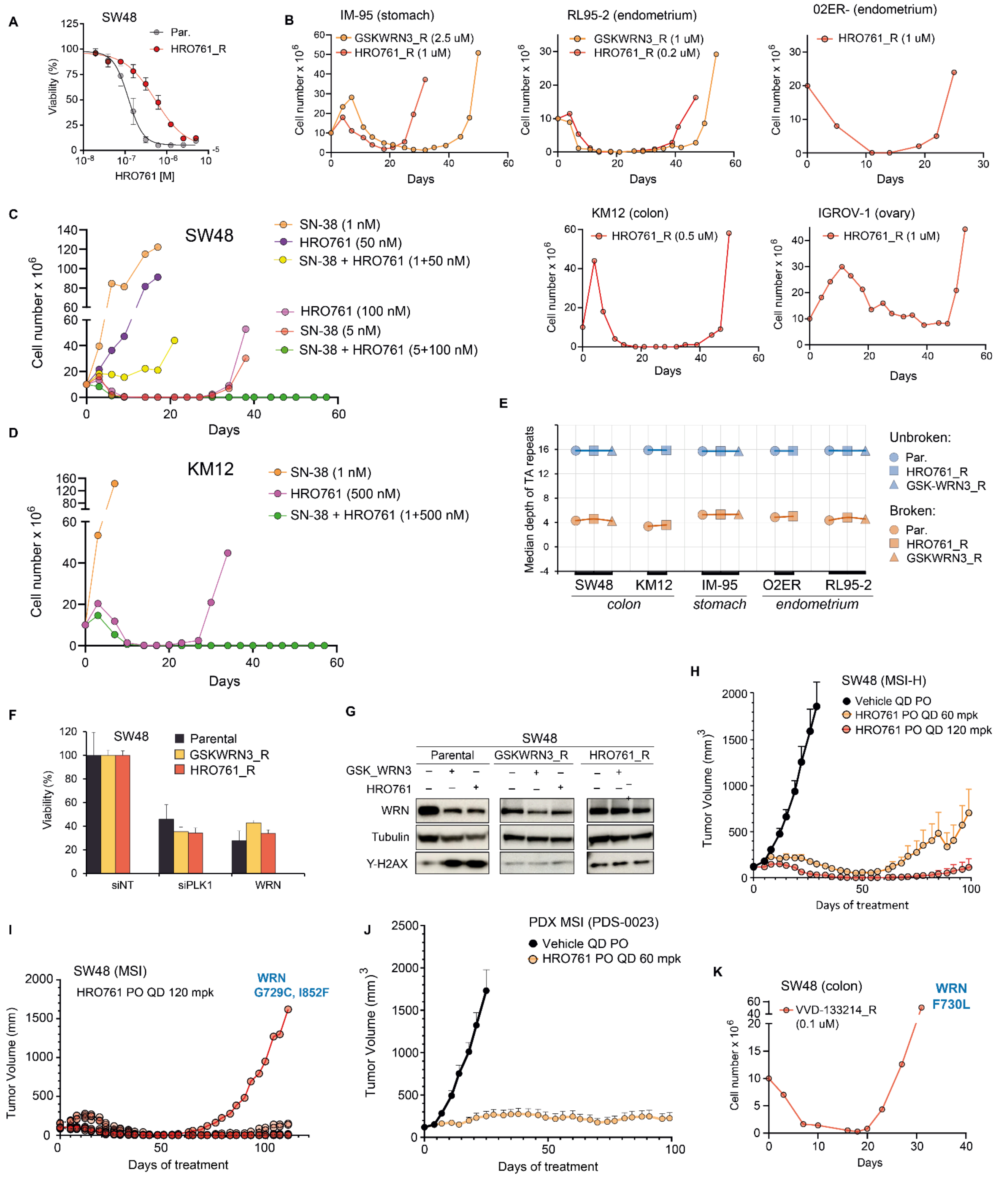


***Supplementary Figure 5. Characterisation of WRNi Resistance in MSI Cancer Cells, Xenografts and Patient-Derived Organoids.*** *(A) Dose–response curves showing reduced sensitivity of HRO761-resistant SW48 cells compared to parental controls. (B) TTP assays in additional MSI cancer cell lines (KM12, RL95-2, 02ER, IM-95) treated with WRN inhibitors GSK_WRN3 and HRO761 reveal consistent resistance profiles, similar to those observed in SW48. The tissue of origin and drug concentration used for selection in each model are indicated in the corresponding sub-panel. (C–D) TTP assays in SW48 (C) and KM12 (D) MSI colorectal cancer cells treated with WRN inhibitors HRO761, SN38 (irinotecan's active metabolite), or their combination. Sublethal and high-dose combinations were tested. Combination treatments delayed growth and the onset of resistance compared to monotherapies. Higher-dose combinations fully suppressed resistance throughout the*

*assay duration. (E) Median sequencing depth at broken and intact (TA)-repeat loci in whole-genome sequencing (WGS) data from parental and WRNi-resistant models across different tissue types. (F) RNAi-mediated WRN knockdown in GSK_WRN3- and HRO761-resistant SW48 cells. Data are mean ± SD of four technical replicates and represent two independent experiments. (G) Immunoblot of WRN protein in parental, GSK_WRN3-resistant, and HRO761-resistant SW48 cells treated with GSK_WRN3 (1 μM) or HRO761 (0.5 μM). Tubulin serves as the loading control. Results are representative of two independent experiments. (H) Average tumor volume for SW48 MSI xenografts treated with HRO761 (60 mg/kg or 120 mg/kg), showing initial regression followed by acquired resistance and tumor regrowth. (I) SW48 MSI tumor xenografts volume measurements for individual mice (n = 10 mice) treated with 120 mpk HRO761. Initial tumor regression is followed by resistance and tumor regrowth in one tumor. The WRN mutation shown was identified post-explant via deep-targeted sequencing, confirming its role in resistance. (J) Tumor volume measurements from an MSI PDX model (PDS-0023) treated with HRO761 (60 mg/kg). Data are the average ± SD of 10 mice per group. (K) The TTP assay in SW48 colorectal cancer cells treated with VVD-133214 monotherapy (0.1 μM) shows a rapid emergence of resistance. WGS highlighted the emergence of the WRN F730L mutation.*

### Supplementary Figure 6

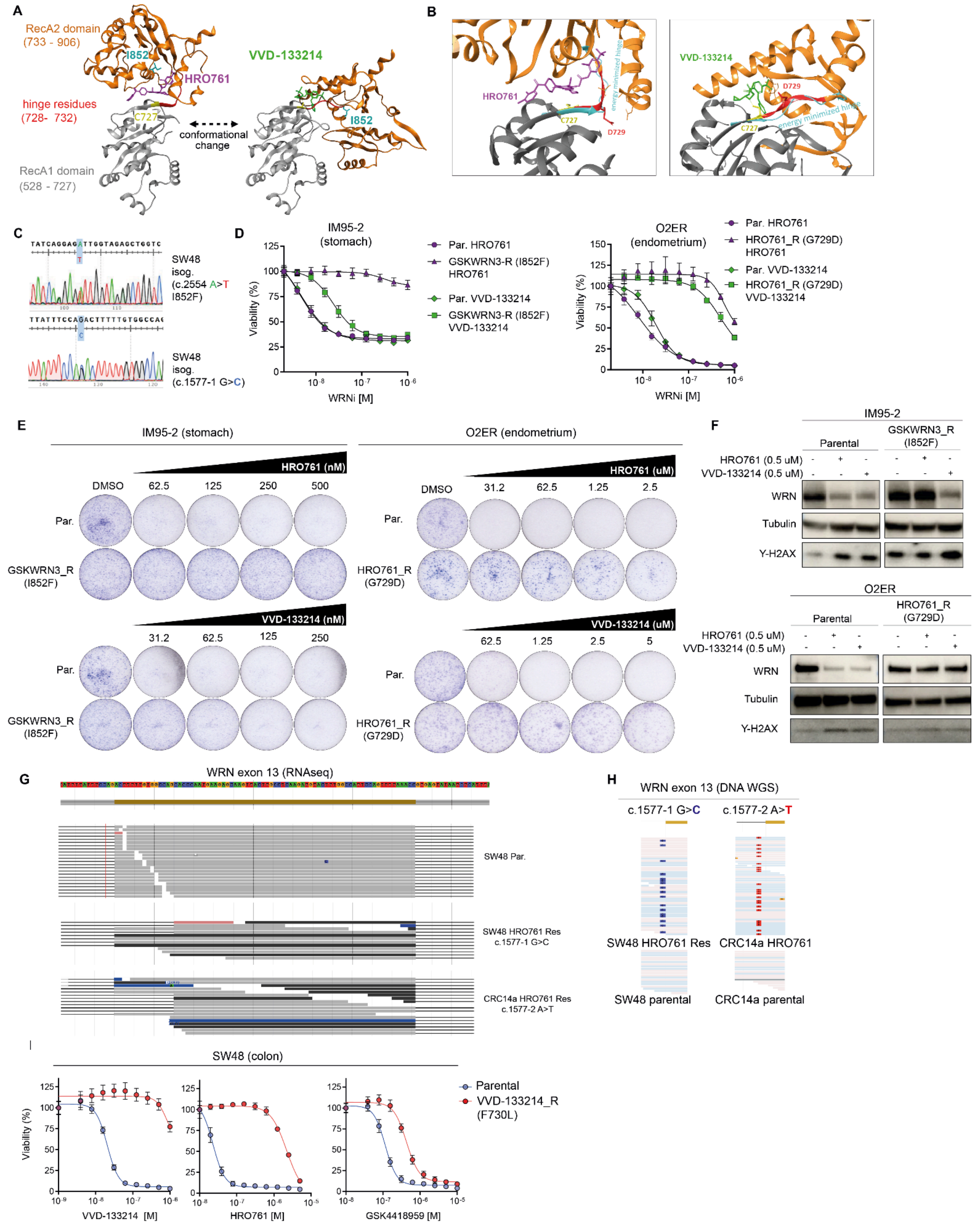


***Supplementary Figure 6. Characterization of WRN Resistance Mutations in MSI Cells and Their Differential Sensitivity to WRN Inhibitors.*** *(A) Structural models of WRN helicase bound to HRO761 (pdb 8PFO; left) and VVD-133214 (pdb 7GQU; right). HRO761 stabilizes WRN in a relatively extended conformation, while VVD-133214 locks WRN in a more compact, inactive form, inhibiting ATPase function. Different elements of the complex are depicted: RecA1 domain residues 528-727 (grey), hinge residues 728-732 (red), RecA2 domain residues 733-906 (orange), C727 (yellow), and I852 (cyan). Ligands HRO761(magenta) and VVD-133214 (green) are shown with sticks. (B) The*

*secondary structure of the hinge with the G729D mutation, after local energy minimization (cyan), showed a different arrangement compared to the initial crystal structures. The surrounding residues around the G729D mutation sites are shown as sticks. (C) Sanger sequencing electropherograms validating the heterozygous introduction of WRN mutations via CRISPR–Cas9 gene editing in SW48 cells. (D) Dose-response viability curves in 02ER (endometrium) and IM95-2 (stomach) cells comparing parental and WRNi-resistant lines (G729D and I852F, respectively) treated with HRO761 and VVD-133214. Data represent the mean ± SD of three technical replicates from three independent experiments. (E) Clonogenic assays in IM95-2 (stomach, left) and 02ER (endometrium, right) and WRNi-resistant lines harboring G729D and I852F mutations, respectively. The Ishikawa (Heraklio) 02ER cell line is referred to as 02ER. (F) Western blot analysis of IM95-2 parental and WRNi-resistant I852F mutant cells treated with HRO761 or VVD-133214, and O2ER parental and WRNi-resistant G729D mutant cells. Blots show WRN degradation and the DNA damage marker γ-H2AX. Tubulin served as a loading control. Representative of two independent experiments. (G) RNA-seq read alignments across WRN exon 13 in CRC14a and SW48 cells. Parental lines show normal inclusion of exon 13, whereas HRO761-resistant clones (CRC14a c.1577-2 A>T and SW48 c.1577-1 G>C) exhibit aberrant splicing, leading to exon 13 skipping. Tracks display read coverage (grey) and splice junctions (black arcs), highlighting the loss of canonical exon 13 inclusion in resistant cells compared to WT. (H) Analysis of WRN exon 13 mutations in WGS. Representative images highlight the mutated residues 1577-1G>C in SW48 and c.1577-2A>T in CRC14a, observed in HRO761-resistant samples. I) Dose–response curves for SW48 parental and VVD-133214-resistant cells (F730L) treated with HRO761, VVD-133214, or GSK4418959.*
